# UV-induced reorganization of 3D genome mediates DNA damage response

**DOI:** 10.1101/2024.05.27.595922

**Authors:** Veysel Oğulcan Kaya, Ogün Adebali

## Abstract

While it is well-established that UV radiation threatens genomic integrity, the precise mechanisms by which cells orchestrate DNA damage response and repair within the context of 3D genome architecture remain unclear. Here, we address this gap by investigating the UV-induced reorganization of the 3D genome and its critical role in mediating damage response. Employing temporal maps of contact matrices and transcriptional profiles, we illustrate the immediate and holistic changes in genome architecture post-irradiation, emphasizing the significance of this reconfiguration for effective DNA repair processes. We demonstrate that UV radiation triggers a comprehensive restructuring of the 3D genome structure at all levels, including loops, topologically associating domains and compartments. Through the analysis of DNA damage and excision repair maps, we uncover a correlation between genome folding, gene regulation, damage formation probability, and repair efficacy. We show that adaptive reorganization of the 3D genome is a key mediator of the damage response, providing new insights into the complex interplay of genomic structure and cellular defense mechanisms against UV-induced damage, thereby advancing our understanding of cellular resilience.

## Introduction

Ultraviolet (UV) radiation, an ever-present environmental hazard, poses a substantial threat to genomic stability and cellular homeostasis. Upon UV exposure, a cascade of events unfolds, including the formation of the most prevalent DNA lesions like cyclobutane-pyrimidine dimers (CPD) and pyrimidine-pyrimidone adducts (6-4PP), capable of disrupting crucial DNA processes [1,2].

Concurrently, UV radiation activates the intricate DNA damage response (DDR) network, spanning cell cycle arrest, inflammation, and apoptosis [3,4]. If not well-orchestrated, it results in genetic mutations, and genome instability, elevating the risk of malignancies and diseases [5,6]. Nucleotide excision repair (NER) stands as the primary mechanism for rectifying UV-induced DNA damage [7,8], and, the development of genome-wide techniques has provided invaluable tools for studying the repair events of UV-induced lesions and assessing the susceptibility of the genome to such bulky adducts [9,10].

Understanding various architectural features of genome structure, including chromosome territories, compartments, and topologically associating domains (TADs), and their temporal dynamics, along with the influence of DNA damage repair mechanisms, is vital to deciphering the DDR machinery. Previous research in the field of radiation-induced DNA damage has primarily focused on the impacts of ionizing radiation on genome structure, particularly X-rays, which delivers a higher energy dose to the chromatin fibres compared to UV. Studies on genome folding have shown that ionizing radiation exposure can cause changes in the spatial organization of the genome upon the formation of chromosomal translocations and double-stranded breaks (DSB). Because of this, most of the experiments evaluated how DSBs cause inflammation and how they are repaired in a 3D genome context [11,12,13,14,15]. It has been reported that, increased TAD strength and insulating efficacy at boundaries occurs after X-ray exposure, and consistent across cell types [16]. As proven by the lack of TAD boundary strengthening observed in ATM-deficient fibroblasts, this suggests an elevation in interaction density due to the ATM kinase’s involvement in modulating the Cohesin complex and facilitating the recruitment of CTCF proteins, in concordance with the TADs span γ-H2AX foci [16,17,18]. Also, efficient repair of DSBs has been reported through chromatin compartmentalization, which mediates the clustering of damaged foci [19]. While these studies have provided crucial insights into the structural changes within the genome following translocation-triggering DSBs, a notable gap persists in understanding how these insights extend to UV-induced DNA damage. Unlike DSBs after ionizing radiation, UV-induced damage primarily results in the formation of pervasive bulky adducts along the genome that needs NER machinery to remove short single-stranded segment that contains the lesion.

Accordingly, we sought to answer the key questions remaining elusive so far for UV radiation: (I) Is UV exposure alone sufficient to induce 3D changes, and does genome folding lead to a more compacted or decompected structure under UV stress?; (II) Do active and inactive compartments of the genome adjust with DDR-mediated transcriptional regulation?; (III) Do the TAD strengthening and boundary insulation correlate with NER efficiency?; (IV) Do the formation or loss of chromatin loops contribute to the facilitation of transcriptional regulation and/or accessibility for NER machinery?; (V) Does the alteration in 3D genome topology play a role in mediating transcriptional regulation for immediate early genes involved in the UV response, such as well-known AP-1 members *JUN* and *FOS*? [20,21,22,23]

Our research endeavors to fill the gaps in understanding and complete this intriguing picture by using both high-resolution contact maps and RNA-Seq to find out how UV radiation affects both the transcriptional regulation and the three-dimensional structure of the genome (Fig. 1 a). We exposed cells to UV radiation (254 nm, 20 J/m^2^) and collected samples at various recovery time points (12 minutes, 30 minutes, and 60 minutes) to comprehensively study the interplay between 3D genome reorganization, gene regulation, and DNA damage and repair. Additionally, we utilized Damage-Seq[10] data collected right after UV exposure and XR-Seq[9] data from 12 minutes later to get a full picture of the dynamic processes that control UV-induced DNA damage and repair. Using single-nucleotide resolution XR-Seq and Damage-Seq maps, we were able to track the temporal changes in DNA repair mechanisms and study the exact locations and types of UV-caused DNA damage.

**Figure 1:**
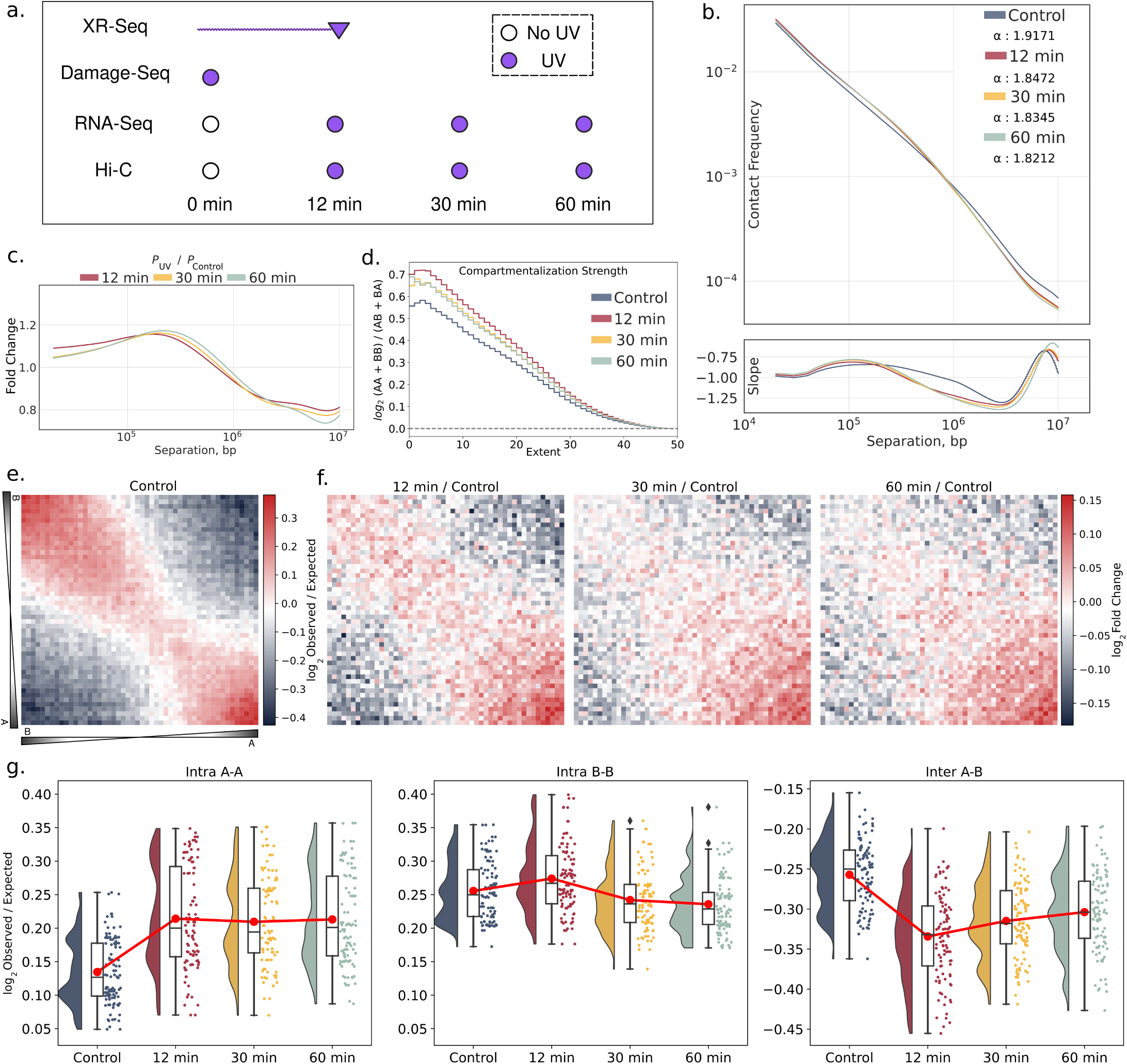
The immediate effect of UV in chromatin conformation and ongoing alterations in the first hour. **a.** Experimental datasets used in the study. For both lesion types (6-4PPs, CPDs), XR-Seq captures the previous repair events, while Damage-Seq captures lesions’ locations on the genome. **b.** Decay plot on 10kb resolution (on log_10_-log_10_ scale) and its slope computed with central differences. Decay parameters (α exponents) of fitted powerlaw distrubutions for each sample. **c.** Decay fold-changes normalized with non-UV control. **d.** Saddle strength of the samples, where the x-axis shows the increasing extent of the compartment percentiles. y-axis show normalized contact strength of (AA+BB) / (AB+BA). (Methods) **e.** Saddle-plot of the non-UV sample (100kb resolution) shows interaction intensity with respect to each compartment percentiles. **f.** Saddle-plot with normalized contact intensities of post-UV samples, relative to the non-UV sample. **g.**. Top 20% compartments normalized interaction intensity comparison. Red connectors point out the means.

Furthermore, we follow a deductive analysis with respect to hierarchical organization of the genome at different scales. Employing a multi-omics and four-dimensional (4D) methodology enabled us to gain a better understanding of how UV radiation affects the 3D organization of the genome and how this affects gene regulation, expression, and DNA repair processes, as well as damage formation in terms of physical and biological relevance. In this study, we show an immediate and holistic response of genome maintenance to UV exposure and ongoing structural reorganization.

## Results

### UV exposure favors short-to-mid range interactions

We conducted Hi-C sequencing before and at 12, 30, and 60 minutes post-UV exposure, ensuring ample sequencing depth for subsequent comparative analysis (Fig. 1 a, Supp. Table1). As calculated (see Methods), all Hi-C samples exhibited sufficient coverage for analysis at a 10 kb resolution. Also, we carried out RNA-Sequencing under identical treatment conditions as Hi-C sequencing, as well as our previously generated XR-Seq and Damage-Seq datasets [24].

One notable observation from the contact maps is the clear decrease in contact frequency as genomic separation increases, a phenomenon often termed distance-dependent decay [25,26]. Such distinctions in characteristics have previously been associated with variances in biological processes, such as differentiation in cell types and cell cycle stages [27]. To understand the underlying changes in the chromosome conformation dynamics, we calculated the contact frequencies with respect to genomic separation (Fig. 1 b). In distance decay analysis, the non-UV sample exhibited different characteristics. A trend is valid for short-to-mid range interactions (< 1Mb), favoring higher frequency for UV-exposed samples. Yet, for long-range (1 Mb ∼ 10 Mb) interactions, non-UV sample propose higher frequency. We computed the slope by employing central differences within each distribution, revealing that the rate of change varies between long, and short-to-mid range interactions. We also fit power-law distributions to obtain α exponents for each sample to show how quickly the decay occurs, and the non-UV sample proposed the largest value, differentiating from the UV-exposed samples, and showing the rapid loss of the tail of the distribution for the non-UV sample. The magnitudes of α values follow the time-course trend, meaning that short-to-mid range interactions are getting stronger by time after UV irradiation.

To better identify the specific ranges of genomic separation in frequency change occurs, we have calculated the fold changes of the distance-dependent decay for each UV-exposed sample compared to the non-UV control (Fig. 1 c). Compared to the non-UV sample, UV radiation promptly favors short-to-mid range interactions in a timely manner. This shows, short-range interactions (<100Kb) are primarily favored in 12 minutes, and the highest frequency of mid range interactions (100Kb ∼ 1Mb) are seen in 30, and 60 minutes. These results might be related to the initial local compaction in the first 12 minutes after the UV and following at the latter time points of 30 and 60 minutes, relatively relieved, yet still more compact chromosome conformation compared to non-UV.

### UV stress strengthens intra-compartment interactions

The genome is partitioned into distinct spatial compartments, such that stretches of active chromatin tend to lie in one compartment, called the A compartment, and stretches of inactive chromatin tend to lie in the other, called the B compartment[25,26,28]. Chromosome regions with similar compartment profiles have a higher tendency to interact with each other. Specifically, active regions have a greater frequency of contact with other active regions, whereas inactive regions are likely to have more frequent interactions with other inactive regions. Accordingly, we have identified compartments with the principal eigenvector of the contact matrix of non-UV sample, by stratifying genomic regions into 50 percentiles with similar values of the eigenvector (Fig. 1 e, see Methods).

Consequently, we assessed the strength of the compartments and determined that the most powerful interactions within the compartments take place 12 minutes after the UV exposure, followed by similar strength between 30, and 60 minutes, in which all of the UV-exposed samples showed high compartment strengths (Fig. 1 d, see Methods). To accompany this finding, we wanted to elucidate which compartments are majorly differentiated with respect to contact strength after UV exposure.

We found that interactions within compartments showed an immediate increase, predominantly within active A compartments, while interactions between A and B compartments showed an immediate decrease at 12 minutes, when compared to non-UV sample (Fig. 1 f). However, this trend is partially followed at 30, and 60 minutes, due to the strength of interactions within B compartments returned to similar levels of non-UV sample. We also evaluated the interaction strength changes at the top 20% compartmental groups, for A and B (Fig. 1 g). Intra A-A interactions post-UV showed the most increase with elevated levels. In contrast, Intra B-B interactions showed an increase at 12 minutes, but followed with reduced levels of interactions at 30, and 60 minutes. Also, a trend can be seen with inter A-B interactions, where levels of interactions decreased after UV exposure, in contrast to immediate increase in intra-compartment interactions.

### Highest dynamicity of compartmentalization is present within transcription regulatory genes

Chromatin compartment profiles and interaction topology within active and inactive compartments are influenced by shifts in gene expression during cell differentiation [29], senescence [30,31,32] or in stress conditions [33,34]. To understand the relationship between compartment profiles and DDR-mediated transcriptional regulation, we sought to perform an exploratory analysis between RNA-Seq and Hi-C datasets.

Accordingly, we performed a correlation analysis between compartment profiles among time points (Fig. 2 a). To pinpoint sets of genes exhibiting varying expression levels over time, we first conducted a time series analysis (referred to as TS analysis afterward) to obtain differentially expressed genes (DEGs) that show significant differences across the time points (Supp. Table 3). We employed expression profile clustering to organize these genes into clusters based on their time-course expression patterns (Supp. Table 4). Each DEG is assigned to a cluster by maximizing the conditional probability, using the maximum a posteriori rule[35] (See Methods). This approach yielded clusters representing different patterns and levels of gene expression changes over time (Fig. 2 b).

**Figure 2:**
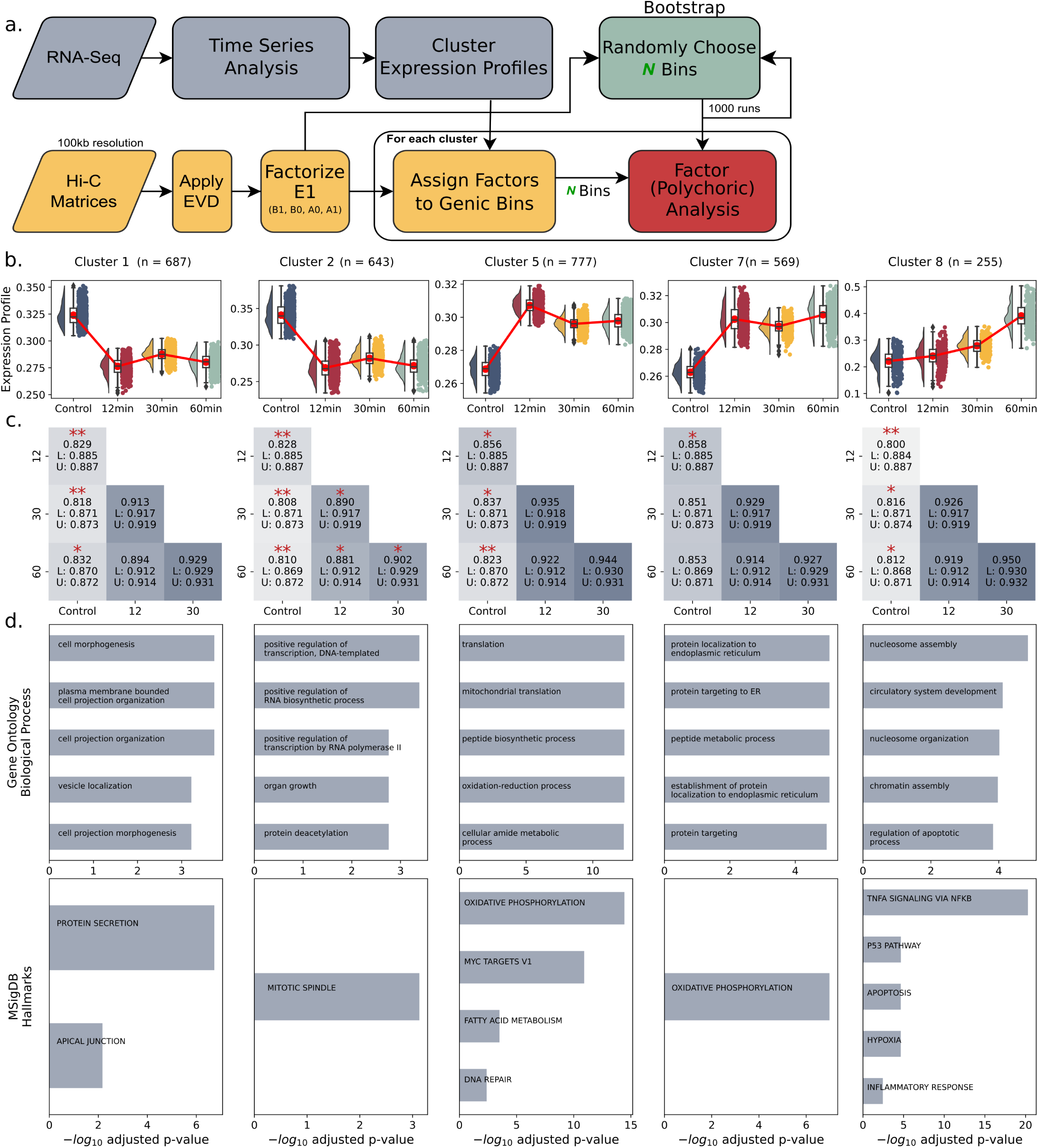
Correlation analysis of compartment profiles under UV stress. **a.** Schematic illustrates the process applying factor correlations on the compartment profiles overlapped with differential gene expression patterns. **b.** Raincloud plots show expression profiles of each DEG cluster with respect to time points. Red connectors point out the means. **c.** Heatmaps show compartment profiles correlation between timepoints per cluster. Asterisk signs show the 2-tailed p-values (* <= 0.1, ** <= 0.01). Lower and upper bounds of confidence interval (95%) of bootstrapped (expected) distribution are presented with L and U. **d.** Barplots show significantly enriched terms (Adjusted p-value < 0.05) for each DEG cluster in GO-BP and MSigDB Hallmarks databases. x-axis shows significance (-log_10_adjusted p-values), y-axis shows the enriched term

Concurrently, we applied eigenvector decomposition to each contact matrix of time points to derive principal eigenvectors, enabling the identification of A and B compartments. We factorized the genomic bins (100kb bins) into factors with similar eigenvector values, denoted as B1, B0, A0, or A1. Each genomic bin, linked to its corresponding factor, was then intersected with genes from each cluster. For each pair of time points, we conducted polychoric factor correlation on the factors associated with the intersected regions. In general, factor analysis showed higher correlations (∼0.90-0.95) within the UV-exposed samples across the three time points, while the correlations remained consistently lower (∼0.80-0.85) for UV-exposed samples compared to the non-UV control sample (Fig. 2 c). This observation suggests that, compartmentalization changes between pre/post-UV are associated with the expression level differences, implying structural reorganizational effect on post-UV gene regulation. To assess the significance of these correlations, we utilized bootstrapping, which involved randomly selecting an equal number of genomic bins multiple times from the dataset. We then performed polychoric factor correlation on these selected regions between the time points. By comparing the observed correlation from the actual data with the distribution of correlations obtained from bootstrapping, we generated empirical p-values. These p-values indicate whether the observed correlation between conditions significantly deviates from the correlation expected based on the bootstrapped distribution.

Furthermore, we have performed gene set over-representation analysis on the DEG clusters to obtain the enriched terms for biological pathways and molecular signatures. Accordingly, we have manually selected 5 clusters primarily based on the presence of enrichment terms for Gene Ontology Biological Process (GO-BP) [36] and The Molecular Signatures DB Hallmarks [37] (MSigDB Hallmarks) (Fig. 2 b, c, d, Supp. Table5). Overall, enrichment of these clusters revealed that under UV stress, genes related to positive transcription regulation are down-regulated, and genes on translation and repair/cell death-related pathways are up-regulated (Fig. 2 b,d). For clusters 1,2,5 and 8, we have observed that factor correlation is significantly lower for comparisons of all of the UV-exposed samples against non-UV while showing a consistent pattern in parallel with the changes in the expression profiles. Remarkably, cluster 2 exhibited notable variability in factor correlations across each time point comparison, implying a dynamic behavior. The gene set involved in positive transcriptional regulation experiences repression immediately after UV exposure, which suggests overall transcriptional repression during DNA damage response.

### Immediate TAD boundary strengthening after UV exposure

A topologically associated domain (TAD) is a genomic segment where DNA sequences have a higher propensity to interact with each other compared to sequences outside the domain, where these domains are demarcated by boundaries [38]. As per the existing theory regarding TAD formation, genome regions are dynamically gathered within TADs via loop extrusion facilitated by the cohesin complex until an insulator protein like CTCF obstructs the extrusion process [39,40,41,42]. However, the potential consequences of UV damage on TAD boundaries remain unknown. To evaluate the impact of UV on TADs, we identified the boundaries of TADs and measured the strength of each boundary, indicating the degree of separation between TADs.

We found that boundary strengthening takes place at each time point after UV exposure compared to the non-UV sample (Fig. 3 a). This shows that the ratio of outgoing interactions from TADs to within TAD interactions is significantly decreasing, proposing a model in which TAD boundaries are more open and accessible with increased insulation in 3D setting. While there is a significant increase (p<1e-4) in boundray strengths between pre/post-UV samples, there is no significant difference in mean boundary strengths among UV-exposed samples (Fig. 3 a). Following this, we revealed the ratio the boundaries are preserved across time points and found that most of the boundaries are preserved across transitions (Fig. 3 b). We also found that a smaller number of boundaries are non-preserved, with a trend that after each transition, the number of non-preserved boundaries decreases, with the highest occurring at the transition to 12 minutes. The term “non-preserved boundaries” refers to regions not identified algorithmically due to their weak boundary characteristics in the tested sample (see Methods).

**Figure 3:**
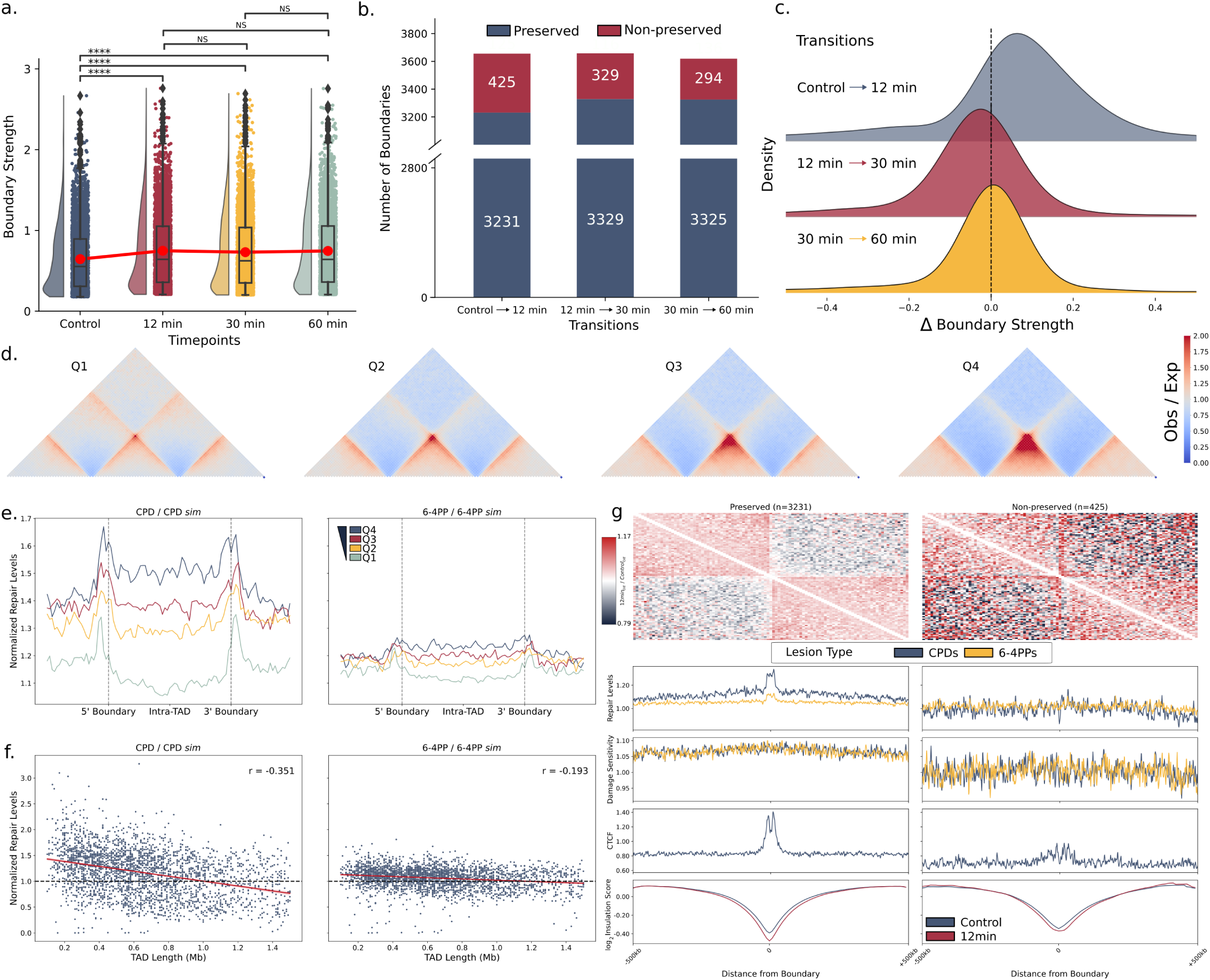
TADs & boundaries show changes post-UV in accordance with repair and damage profiles. **a.** Mean boundary strength is increased post-UV, with 12 minutes being the highest. (Mann-Whitney U-test, **** means p-value <= 1e-04) Red line connectors point out the means. **b.** Number of boundary states per transition across time points. **c.** Ridgeline plots show a comparison of boundary strength changes per preserved boundary set at each transition. **d.** TADs were grouped by their strengths into quartiles. **e.** Normalized repair levels are plotted with respect to TADs grouped by their strengths, CPDs (Left), and 6-4PPs (Right). **f.** Normalized repair levels are plotted with respect to TADs sizes, CPDs (Left) and 6-4PPs (Right), Pearson correlation p-values < 0.001. **g.** The top row shows a pile-up analysis of aggregated and averaged on-diagonal snippets centered at boundaries. The bottom rows show normalized repair levels (XR-Seq) and damage sensitivities (Damage-Seq) of both lesion types, CTCF ChIP-Seq (nontreated HeLa-S3) fold over input data, and insulation scores. Columns are separated with respect to preserved and non-preserved boundaries of the first 12 minutes of transition post-UV.

To better characterize these transitions, we calculated the change in the boundary strength of the preserved boundaries. Among the transitions between the time points, the largest positive change was observed right after the UV exposure, in the first 12 minutes (Fig. 3 c). The transition from 12 to 30 minutes showed a slight decrease in boundary strengths, followed by a relative consistency at the 30 to 60-minute transition. This immediate effect proposes that, similar to boundary preservation, the highest boundary strengthening response is seen in early post-UV. Between 30 and 60 minutes, the boundary strengths remained consistent.

### Eficient UV repair in TADs is correlated with boundary strength

Since the boundaries are proposed to be more open in 3D setting with respect to boundary strength after UV exposure, we aimed to investigate repair in TADs to see whether such characteristic is favored for normalized repair levels of CPDs and 6-4PPs, on the regions within TADs, boundaries and flanking regions. We scored TADs as the ratio of two numbers, within-TAD intensity to between-TAD intensity, as previously described [43]. We grouped TADs into quartiles with respect to their strengths (Fig. 3 d) and projected normalized and averaged repair levels at these regions (Fig. 3 e). Overall, boundaries showed enriched repair levels. For both CPDs and 6-4PPs, the highest repair levels are associated with the TADs that are in the top quartile (Q4), which also have enriched intra-TAD repair levels compared to flanking regions. Interestingly, as the TAD strength decreases, intra-TAD repair levels decrease compared to flanks (Q2 and Q1). Relative to 6-4PP, CPD repair is more affected by the TAD boundary strength, which in line with the fact that CPD repair efficiency is relatively more prone to chromatin factors [10,44]. We questioned whether repair levels change with the increasing TAD size. In general, average repair levels of both damage types show a negative correlation, with CPDs showing a higher negative correlation (r: -0.351, p-val: 3.518e-82) compared to 6-4PPs (r: -0.193, p-val: 6.947e-25) (Fig. 3 f). Accordingly, we speculate that the negative correlation between repair levels and increasing TAD size might be related to the repair events initiating at the boundaries and possibly lowered the probability of accessibility for repair elements along a longer TAD.

### Loose boundaries have low UV damage sensitivity and repair

To see if boundaries provide an accessibility zone for repair elements, we wanted to compare damage response in the preserved and non-preserved boundaries. To understand the interaction change surrounding boundaries, we averaged the on-diagonal pile-ups centering the boundaries, comparing contact maps of 12 minutes to non-UV (Fig. 3 g). It is clear that TADs spanning both preserved and non-preserved boundaries have elevated within-TAD intensities. However, between-TAD intensities decline relative to within-TAD intensities for TADs spanning preserved boundaries, providing higher accessibility to boundaries. For TADs spanning non-preserved boundaries, we see an elevated intensity of the interactions between TAD, resulting in a less accessible zone for non-preserved boundary regions and possibly leading to a structure where adjacent TADs merge. Accordingly, we have obtained publicly available fold-over-input data for nontreated HeLa-S3 CTCF ChIP-Seq (ENCSR000AOA) to compare occupancy at boundaries. Non-preserved boundary regions showed lower levels of CTCF occupancy at minute 0, proposing a lower potential to insulate TADs and after under UV radiation, possibly leading to acute loss of boundary characteristics for certain TADs.

Interestingly, TADs spanning preserved boundaries showed higher damage sensitivity. Furthermore, we wanted to understand the contrast in the repair levels between the boundary states. Overall, we have observed a lower efficiency in repairing lesions at non-preserved boundary sites. In contrast, elevated repair levels have been noted for both CPDs and 6-4PPs at the preserved boundaries, with CPDs exhibiting a particularly higher efficiency. Thus, we observe that higher insulation efficiency of strong boundaries post-UV correlates with greater repair efficiency, in contrast to their weaker counterparts.

### CTCF-dependent loops are enriched post-UV

Enriched contact frequency peaks that appear as distinct points in mammalian contact maps are a common characteristic, and they propose significant interactions [26]. These points represent ‘loops’, and may manifest individually or as components of grids, often found at domain corners. Accordingly, chromatin loops are considered the primary mechanism of enhancer-promoter interactions facilitated upon cohesin-mediated loop extrusion accompanied with barrier elements [45,46,47]. A chromatin loop consists of a pair of proximal regions, referred as loop anchors. For all time points, we have identified chromatin loops and created the intersection sets for loops and their anchors (Fig. 4 a). We have observed that the number of anchors common in all samples (51.0%) exceeds the proportion of loops common in all samples (31.6%), indicating that certain shared anchors are involved in forming different loops over time after UV-exposure.

**Figure 4:**
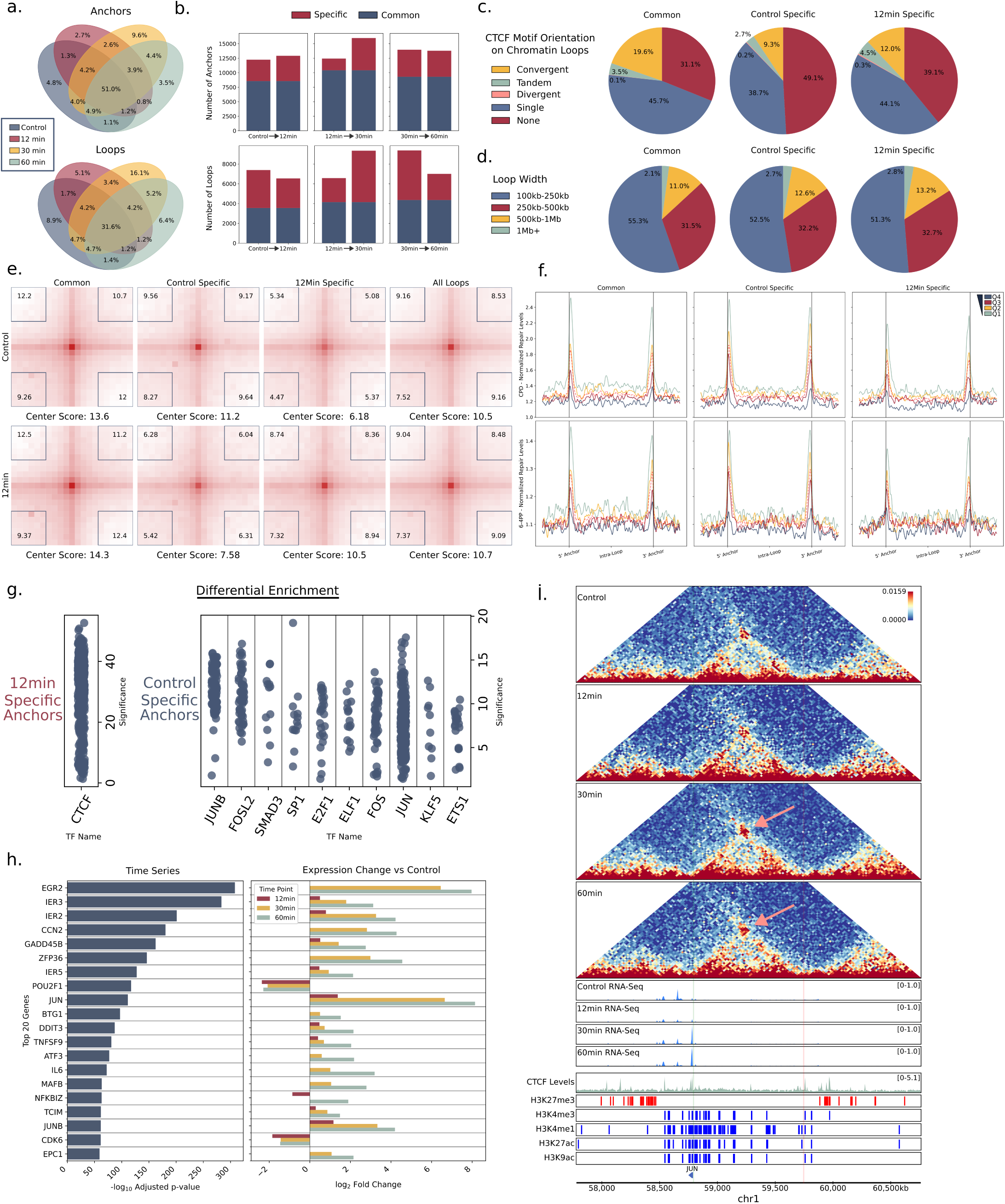
Loops & anchors show differential characteristics post-UV. **a.** Venn diagrams show the percentages of common and specific anchors and loops. **b.** Stacked bar plots show the number of common and specific anchors for each transition. **c.** For the non-UV and 12-minute common and specific anchors, pie charts show CTCF presence and orientation, and **d.** loop size distributions across size ranges. **e.** For the non-UV and 12-minute common and specific anchors, off-diagonal pile-up analysis shows intensity change centered at the loop anchors. Corner numbers correspond to center-to-corner ratios. **f.** Loops are grouped into quartiles based on loop strengths (Methods). Repair levels (y-axis) are plotted along flanking regions and loop regions. Red dashed lines show average repair levels in the associated set. **g.** Unibind differential enrichment analysis with respect to specific anchors of non-UV Control sample, and 12 minutes, separately. Significance shows -log_10_ p-values. **h.** Top 20 genes exhibiting significant changes in their time-course expression profiles with differential looping patterns. **i.** Grouped Hi-C and RNA-Seq genome tracks, with CTCF, and histone markers for JUN locus. Blue and red colors for histone markers signify activation and repression function, respectively. Only JUN is written on genes track for simplicity. Green vertical line shows JUN TSS, and red vertical line shows the corresponding looping region.

For each timepoint transition, we separated loops and anchors into two categories: common and specific loops and anchors. The separation tolerates a range for a ±50kb, meaning neighboring regions are taken into account when determining overlap. Specific loops to a time point indicate a gain or loss of contact strength relative to the transition, as identified by their significant enrichment in only one of the contact matrices (see Methods). Accordingly, we wanted to understand the number of loop/anchor differences between transitions (Fig. 4 b). After UV exposure, we have seen the number of loops has decreased at 12 minutes. At 30 minutes, we have seen the highest number of specific anchors, leading to the highest number of specific loops. This phenomenon did not continue with 60 minutes, by losing most of the specific loops that had been acquired at 30 minutes. We have also seen the number of common loops increase with respect to each transition post-UV.

Hereinafter, we have concentrated on the comparison of non-UV sample and the first time point of 12 minutes to see the primary effects of UV exposure on loops. CTCF is the predominant protein concentrated at loop anchors when arranged in a convergent orientation [48]. Consequently, we aimed to understand the paired orientation of CTCF (Fig. 4 c), while also considering the presence of single and non-CTCF motifs on the corresponding loop anchors. The initial observation was that common loops had anchors with a higher percentage of convergent orientation, which is the most favored orientation for paired motifs. Also, we have seen that loop anchors that are specific to the non-UV Control sample do not possess the CTCF binding motif, with the highest percentage of 49.1% among the three groups, which declined to 39.1% after UV exposure. This implies that certain transitory loops are lost under stress conditions, and loops are formed through CTCF, among other barrier elements, when exposed to UV. Furthermore, we did not detect significant differences in loop widths among the 303kb, 325kb, and 320kb categories for common, control-specific, and 12min-specific loops (medians of 230kb, 230kb, and 240kb, respectively), nor did we observe any substantial differences in composition (Fig. 4 d).

With pile-up analysis, we have compared normalized enrichment by averaging off-diagonal contact matrix snippets centered at loop anchors (Fig. 4 e). We have seen clear enrichment gain and loss after UV exposure to specific loops. Enrichment scores of pile-ups are calculated with the corner scores (center-to-corner ratios).

We further focused on understanding repair levels at specific loops (Fig. 4 f). To do so, we grouped specific loops into quartiles based on their loop strength scores (see Methods). In all comparison groups, we consistently observed that loop anchors exhibited higher repair activity compared to the flanks and the loop body. Additionally, we noted that loops with the lowest strength displayed higher normalized repair levels across all groups. Also, specific loops in the non-UV Control sample, meaning loops that decreased enrichment after UV, showed higher repair levels for both lesion types. However, loops formed post-UV through loop extrusion (12min specific) showed lower repair levels in general.

To identify potential factors of transcriptional regulation with respect to specific loop anchors for non-UV and 12 minutes post-UV, we performed differential enrichment at these regions using transcription factor (TF) ChIP-Seq data available on the Unibind database [49] (Fig. 4 g). For anchors specific to non-UV Control sample, indicating loops that lost strength after UV exposure, we see a significant enrichment for TFs, such as the JUN family, a well-known protein for response to UV radiation, increasing cells’ survival capabilities. Besides JUN, the enriched TFs of the non-UV control-specific anchor set were enriched with FOS and FOSL2, members of the AP-1 transcription factor. This suggests that the potential binding of these TFs is correlated with a decrease in loop strength and higher repair levels. On the other hand, specific anchors to 12 minutes showed significance only for potential CTCF binding, suggesting that the loops are formed through loop extrusion, which is in line with the Fig. 4 c.

Furthermore, we aimed to investigate whether the transcriptional regulation of genes exhibiting significant changes in expression over time was associated with differential looping. Accordingly, we revisited time series analysis (TS), and identified the top 20 genes based on TS analysis significance, that either gained or lost a new chromatin loop at any of the timepoints. We collected the loop anchors containing the transcription start sites (TSS) for the genes of interest (canonical transcript per gene selected with the MANE project[50]) for each time point. Accordingly, we have analyzed differential looping patterns to better characterize expression changes (Fig. 4 h). We observed that complementary looping patterns associated with changes in expression were present for the selected genes. For instance, in the case of the UV-induced immediate early gene JUN, it shows significantly high expression levels at 30 and 60 minutes post-UV. Correspondingly, as pointed on the Hi-C matrices, there is a progressively increasing interaction intensity, particularly notable at 30 and 60 minutes post-UV. In this interaction, one anchor involves JUN, while the other anchor corresponds to an enhancer region with an activation function (Fig. 4 i). The facilitative role of the 3D genome in transcriptional regulation is also relevant in this context following UV exposure.

### Graph neural network segregates genome with respect to 3D dynamics

Our objective was to investigate the facilitative aspects of 3D genome dynamics. To do so, we centered our investigation on the transition from the non-UV control to 12 minutes, using it as a pivotal reference point. This approach enabled us to gain deeper insights into the impact of immediate early genes post-UV exposure and to better align our analysis with repair and damage maps. Up to this point, our analysis has primarily been hierarchical and exploratory, emphasizing genome architectures identified through predefined rules. However, recognizing the highly dynamic nature of genome folding, especially under genotoxic conditions, we acknowledge the necessity of studying regions that exhibit differentiation in higher-order dimensions, rather than relying solely on monotonic metrics like insulation scores or a set of kernels. As a result, we have chosen to leverage Graph Neural Network (GNN) methodology to search into topological alterations between pre-UV and post-UV states, with a particular focus on the initial repair response.

GNNs have been started to utilized in the context of Hi-C, such as for multi-omics integration [51,52,53], 3D genome reconstruction [54], contact matrix imputation [55], identifying differential point-wise interactions [56], and for the double-stranded break (DSB) prediction [57]. Accordingly, we’re utilizing graph representation learning to exploit the inherent relational characteristics of Hi-C interactions, which are effectively captured in a graph structure.

In essence, our task involves determining whether a graph derived from one contact matrix (referred to as the “query graph”) is preserved within another graph (referred to as the “target graph”) generated from a different contact matrix. We aim to identify regions (genomic coordinates) centering the topology change by quantifying deviations in subgraph isomorphism using a GNN (Fig. 5 a). This involves measuring the extent of violation (topology change) of a query graph compared to its paired target graph [58]. Accordingly, we aim to segregate genomic bins into distinct clusters based on topology change characteristics (Fig. 5 c) and gain insights into the intersection of 3D genome alterations and DNA damage/repair processes.

**Figure 5:**
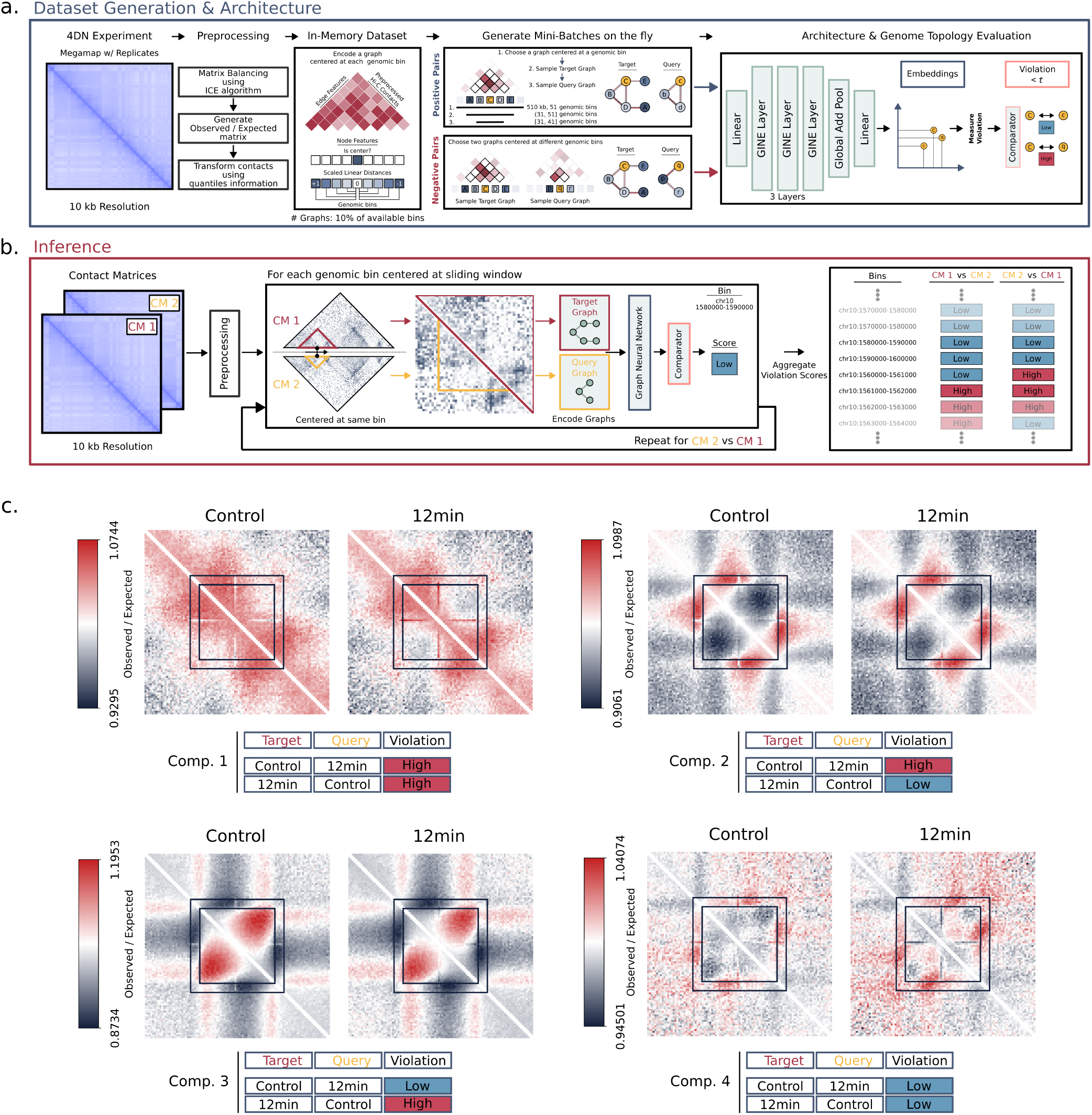
Analysis of genome folding with GNN shows different topologies post-UV. **a.** Schematic illustrating the workflow for the preprocessing steps of contact matrices, dataset generation/graph encoding, and generating different 3D genome topologies to create a violation landscape for GNN training. ** **b.** Schematic illustrating the inference workflow for the violation scores (topology change) from two contact matrices to be compared using the sliding window operation. **c.** With violation scores (Low or High) and the direction of the comparison, the genome is segregated into four groups. Central coordinates in the on-diagonal pile-ups are scored regions. Pile-ups show ± 500kb flanking regions. Black square patches show target (±250kb) and query (±200kb) regions for the generated graphs, respectively.

To create a violation landscape for the GNN’s learning process by simulating varying levels of 3D genome topology changes, we preprocessed data from the 4D Nucleome megamap of merged replicates of non-treated HeLa-S3 cells, and curated a graph dataset. (Fig. 5 a, Methods). Within the graph dataset, the Hi-C contacts are represented as edge features, while the central node and linear distances are encoded as node features. To facilitate the training of the subgraph prediction function aimed at discerning differentiation in genome topology, we created both positive and negative pairs consisting of target and query graphs. Positive pairs consist of target and query graphs centered on the same genomic bin, while negative pairs are sampled from two distinct graphs centered on different genomic bins. This sampling approach ensures lower violation scores for similar topologies and higher violation scores for different topologies. This process involves training a 3-layered Graph Neural Network (GNN) to derive graph-level embeddings for both the target and corresponding query graphs. These embeddings are subsequently utilized to compute violation scores (magnitude of topology change) for each pair of target and query graphs. Following that, these scores serve as input to train a comparator, which learns to discern between low and high violation score labels by determining an appropriate threshold.

To infer changes in genome topology between non-UV and 12 minutes post-UV, we preprocess the contact matrices and employ a sliding window operation akin to insulation score calculation (Fig. 5 b). At each window (genomic bin), we generated larger target and smaller query graphs central to the same genomic bin from contact matrices to compare. These graphs are constructed using the adjacencies and interaction intensities within the sliding window and the same encoding methodology as in the training phase. We measured the violation of queries to corresponding targets with the embeddings obtained from GNN and aggregated violation scores genome-wide. By employing this approach, we capture various topological changes, including both the gain or loss of contacts, as well as differences in the strength levels of preserved contacts (Fig. 5 c).

Subsequently, we categorize genomic bins (10kb resolution) into four groups, referred to as “comparisons,” based on the level (either Low or High) and direction of the violation observed between non-UV and 12 minutes post-UV. With this methodology, we evaluated 88.28% of the genome, excluding blacklisted, balance-masked bins and sparsely interacting regions. The proportions of bins included in comparisons 1, 2, 3, and 4 were 19.79%, 15.15%, 12.35%, and 40.99%, respectively. Accordingly, we merged consecutive genomic bins with the same comparison group label and used central coordinates afterward. For each comparison, we computed the average of on-diagonal snippets from the corresponding contact matrix, thus assessing the normalized contact intensity around the central genomic bin (Fig. 5 c). Initially, we observed that Comparison 4, the largest group with low violation scores in both directions between non-UV and 12 minutes post-UV, exhibited the least dynamicity observed over expected contact intensities. Comparison 1 suggested open and possibly accessible regions in 3D setting, between highly intense short-range interactions in which a high level of violation is present for both directions of the comparison. Between Comparisons 2 and 3, we have seen an interesting contrast between aggregated snippets. In the central regions of Comparison 2, we observed below-expected interactions for closer linear distances to the center, while the surrounding areas exhibited above-expected intensities. On the contrary, Comparison 3 displayed an inverse trend, characterized by above-expected intensities for closer linear distances to the center and below-expected intensities in the surrounding regions. This observation suggests that the direction of comparison, coupled with the differentiation between Low and High levels of violation scores, contributed to a sense of relative symmetry in the analysis while also delineating distinct architectural features.

### Genome undergoes dynamic and relative convulsions favoring repair eficiency post-UV

We aimed to understand the difference in contact intensities within the comparisons in accordance with repair efficiency and damage profiles. Repair efficiency is normalized repair levels with damage sensitivity and respective nucleotide distribution frequency simulations (see Methods). Accordingly, we divided the 12-minute pile-ups into non-UV, generating divide-ups to compare time points at each transition. Again, divide-ups of 4 comparisons yielded different characteristics, in line with violation scores and direction of comparison (Fig. 6 a). In Comparison 1, at the first transition post-UV, central regions exhibited an increase in contact strength with surrounding regions while simultaneously experiencing a decrease in strength for contacts traversing these regions. This characterization was accompanied by high repair efficiency for CPDs being relatively enriched in the evaluated region and were also high on the flanking sites. This favors the hypothesis that these regions bring surrounding regions into close proximity while being accessible themselves, proposing an open and hub-like profile in 3D setting post-UV. On the other hand, Comparison 4, which had the lowest violation scores indicating minimal change in genome topology, showed the lowest repair efficiency.

**Figure 6:**
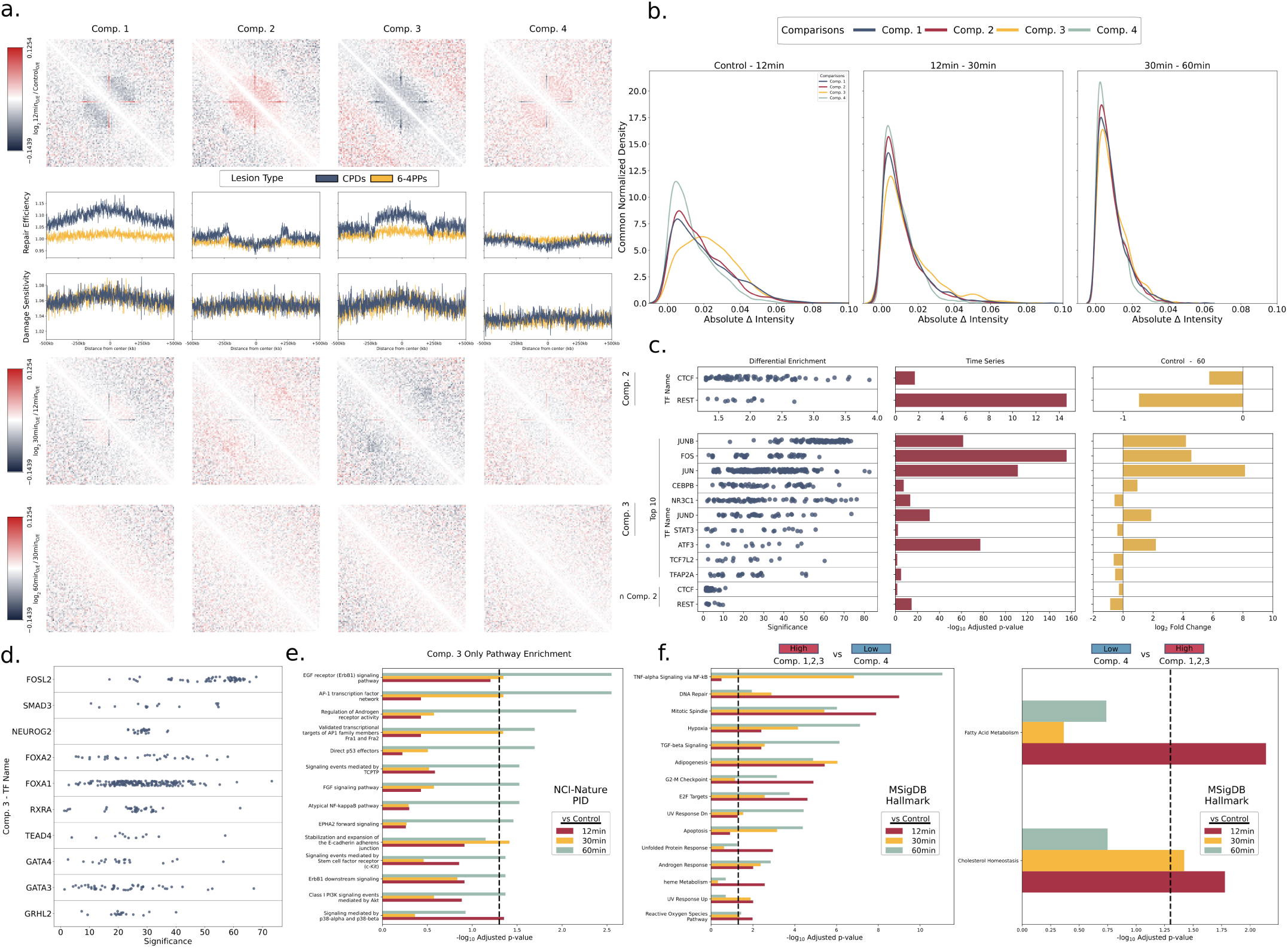
Analysis of genome folding shows differential topologies post-UV. **a.** Top row shows divide-ups of 12 minutes to the non-UV sample. The bottom rows show repair and damage profiles, followed by divide-ups of 30 to 12 and 60 to 30-minute transitions. **b.** Density plots show the absolute intensity changes on the pixel distributions where the pixels’ are located in the target graph region of the corresponding divide-up. Common normalization ensures each area under corresponding density sums to 1. **c.** Unibind differential enrichment analysis for the central sites of Comp. 3 and Comp. 2 for the TFs that their expression is significantly changed. TFs are ordered with respect to the mean significance of the corresponding experiment sets. Significance shows -log_10_ p-values. **d.** Unibind differential enrichment analysis for the central sites of Comp. 3 for the TFs that their expression is stable across time points. **e.** Gene set overrepresentation analysis conducted on Comp. 3 associated DEGS with respect to time points reveals the significance of pathways in a timely manner. Dashed lines show the significance cutoff (p.adj < 0.05). **f.** Unique and enriched terms for High genome topology change (Left), and Low genome topology change (Right). Dashed lines show significance cutoff (p.adj < 0.05)

**Figure 7:**
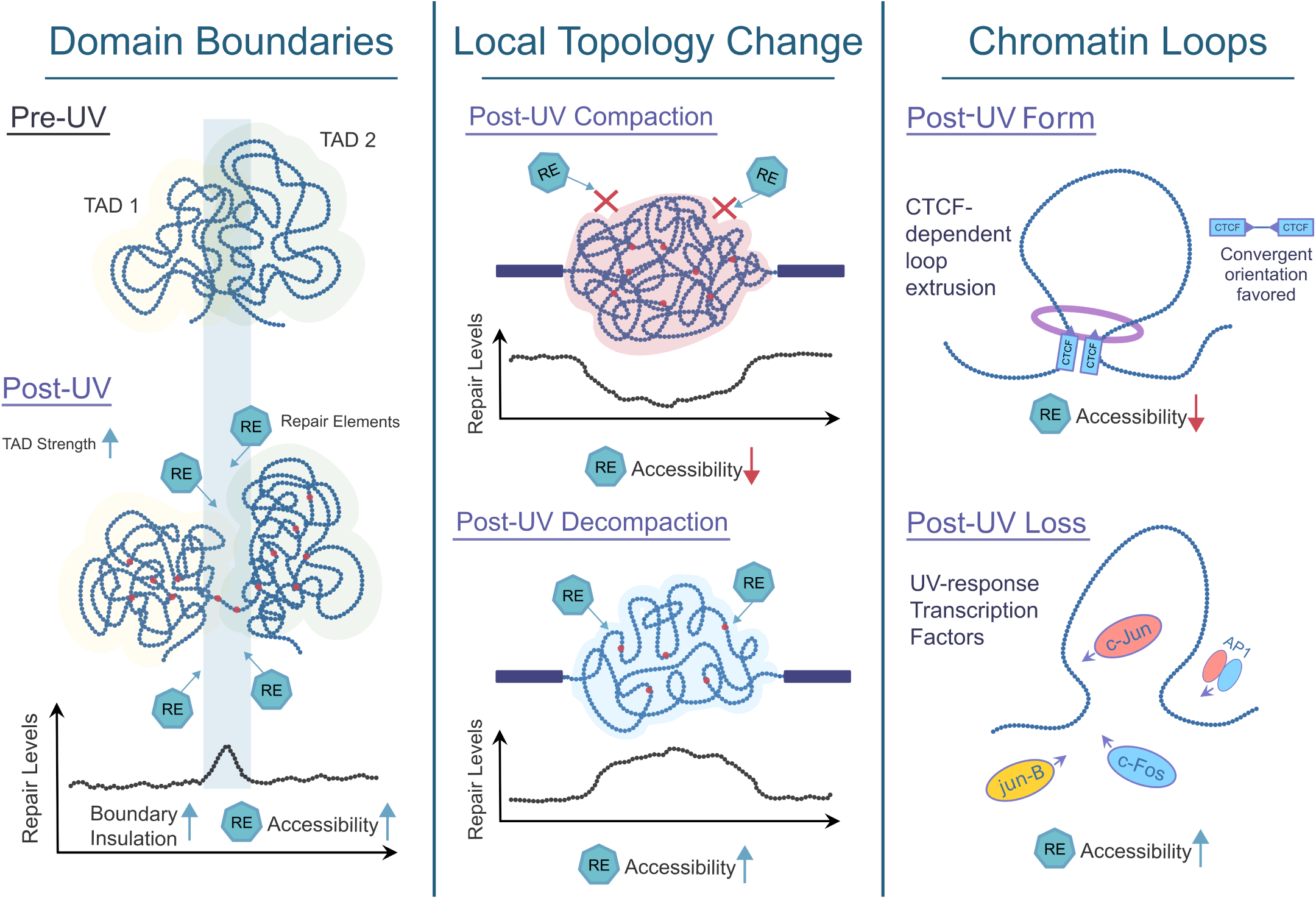
Post-UV 3D genome facilitates UV-induced DNA damage response.

Additionally, we have noted symmetry in the division patterns for Comparisons 2 and 3 and in the repair profiles. We observed that in Comparison 2, central regions with intensities below the expected level demonstrated an increase in strength for both pairwise and surrounding interactions. This observation suggests a transformation into a compact hub following UV exposure. Conversely, in Comparison 3, central regions with intensities above the expected level demonstrated decreased strength for pairwise and surrounding interactions. Furthermore, profiles of repair efficiency for flanking regions exhibited differences between Comparisons 2 and 3, albeit with a notable degree of symmetry within the evaluated regions. Notably, the repair of both lesions exhibited symmetry, which is particularly evident in CPD repair. Additionally, in Comparison 4, we noted the lowest sensitivity to damage, a finding markedly distinct from the other three comparisons. This observation suggests that lower levels of damage correspond to reduced differentiation in genome topology.

In any of the comparisons, we observed a small amount of relative recovery in pixel intensity at the transition from 12 minutes to 30 minutes post-UV exposure. However, we did not observe any discernible difference in the transition from 30 minutes to 60 minutes post-UV.

We also sought to assess the absolute changes in pixel intensity on the divide-ups. We have collected pixel intensities within the GNN evaluated regions on divide-ups, obtained distributions of fold changes on log_2_ scale without direction, and performed kernel density estimations (Fig. 6 b).

Remarkably, we observed that Comparison 4, characterized by the lowest violation scores (lowest topology change), exhibited the lowest pixel intensity changes, favoring the GNN algorithm’s expressive capabilities. We have also observed Comparison 3 is associated with the highest intensity change.

We were especially intrigued by the symmetry observed between Comparisons 2 and 3. Since, in these regions, contrasting 3D genome topologies were associated with compaction and decompaction, respectively for lower and higher repair efficiency. Consequently, we investigated transcription factor binding and leveraged the Unibind experimental data for differential enrichment analysis. We retained only those transcription factors (TFs) whose expression changes were significant in the TS analysis and kept TFs supported by at least 10 individual experiments (Fig. 6 c). Our analysis revealed that decompacted regions evaluated within Comparison 3 exhibited remarkably high levels of enrichment for JUN-family members, with JUNB being highest in mean significance and FOS, and ATF3, indicative of AP-1 member enrichment [59,60,61]. These TFs displayed very high significance levels in the TS analysis, with a notable increase in expression fold-change after the first-hour post-UV. Conversely, compacting regions evaluated within Comparison 2 displayed low but significant enrichment levels for only CTCF and REST. We have not seen any other TF enrichment at Comparison 2 sites, and we also observed significant enrichment of CTCF and REST in Comparison 3. We have also evaluated the TFs that are insignificant in the TS analysis (Fig. 6 d). The most significant enrichment was FOSL2, which is another AP-1 member. Overall, we have seen lower significance levels than the TFs that are significant in TS analysis (Fig. 6 c,d).

Furthermore, we sought to explore how the enrichment of pathways associated with the differential gene expression correlates with 3D characteristics of comparisons (Supp. Table7). To accomplish this, we individually identified differentially expressed genes (DEGs) in the analysis between 12 minutes to non-UV, 30 minutes to non-UV, and 60 minutes to non-UV. We overlapped the bins containing TSS regions of the DEGs in comparison groups separately and conducted gene set over-representation analysis. Accordingly, we wanted to understand signalling pathways that are enriched in Comparison 3, so we included the NCI-Nature Pathway Interaction Database in the analysis [62]. We have observed that “AP-1 transcription network” was significantly enriched, especially for 60 minutes mark post-UV, and included DEGs such as down-regulated NR3C1, and up-regulated FOSL1, FOSB, ATF3, and JUNB (Fig. 6 e). We have also seen that the “UV Response Down” term, which implies genes that are down-regulated post-UV, was only enriched with the DEGs in Comparison 3 and “UV Response Up” only in Comparison 1. Interestingly, we have identified “EGF receptor (ErbB1) signaling pathway”, including down-regulated EGFR, and STAT3 (Fig. 6 e).

Additionally, we wanted to understand the enriched pathways in the comparisons with the high genome topology change (Comparison 1,2,3 combined) in contrast to the low genome topology change (Comparison 4), so we kept only the unique enrichment terms (Fig. 6 f, Supp. Table8). For the genes belonging to the high genome topology change, we have observed enriched terms such as “TNF-alpha signaling via NF-κB” being the most significant, “TGF-beta Signalling” and “Apoptosis” at the 60-minute mark, and “DNA Repair” with the most enrichment at the initial 12 minutes post-UV. However, we have not seen any related hallmarks within the enriched terms unique to low genome topology change.

## Discussion

Recent efforts demonstrated the maintenance of genomic stability within the framework of higher-order chromatin organization [11,12,13,14,16,19]. Indeed, ionizing radiation and other genotoxic agents that primarily form DBSs and trigger translocations cause the 3D genome to play a role in DNA repair mechanisms. Understanding the facilitative role of the 3D genome in efficiently excising bulky adducts is crucial, particularly within the context of our cellular resilience against the most prevalent form of damaging radiation in the electromagnetic spectrum: UV radiation. However, post-UV organizational changes in the 3D genome and its interplay with nucleotide excision repair (NER) and transcriptional regulation have not been studied.

In this study, our focus was particularly on the earliest time points following UV radiation exposure to better characterize the interplay between DNA damage response and the 3D genome. Accordingly, we have generated the contact maps upon UV exposure and performed hierarchical 3D genome analysis. We have demonstrated that UV radiation can induce alterations across all levels of higher-order chromatin organization, initiating an ongoing structural reorganization with certain characteristics that are reminiscent of those seen post-IR. We also observed decreased distal interactions following UV exposure, consistent with findings observed after X-ray irradiation across various cell types, including fibroblasts [16]. Concurrently, we observed an elevation in interaction intensity within compartments and a decrease in inter-compartment interactions following UV exposure, similar to the response seen in DSB-related scenarios across different cell types [14,16]. The loss of distal interactions and increased compartmentalization strength may be potential indicators of a generic response to radiation-induced genotoxic stress.

In our investigation of TADs, we noted a strengthening of boundaries following UV exposure, similar to increased domain insulation seen in X-ray irradiation on different cell types [16]. However, there was also a minor loss of boundaries post-UV. We hypothesize that this loss may be attributed to a physical phenomenon wherein UV radiation induces resonance within the TADs, causing vibration and potentially straining boundary endurance, particularly regarding insulation capabilities. Additionally, we observed increased levels of UV-induced photoproducts in the preserved boundaries, which are likely to be located in highly organized structures where there is little room for physical vibration. Disabled physical elasticity may have led to higher levels of UV energy-induced DNA lesions, such as CPD and 6-4PP. It is plausible to infer that boundaries that can be lost are more resistant to damage possibly because of their less organized structures and more room for absorbing the UV energy.

However, those DNA lesions cannot be efficiently repaired. Conversely, if boundaries are preserved and TADs are better insulated, higher damage rates may occur, yet they are more efficiently repaired. Notably, boundaries emerge as crucial sites as they become more accessible, facilitating repair processes.

The findings prompted us to delve deeper into how higher-order chromatin organization influences the accessibility of the NER machinery. Consequently, we propose a model suggesting that local decompaction of the 3D genome enhances accessibility, potentially easing the recognition and excision of lesions. This model is supported by multiple lines of evidence spanning various levels of 3D genome organization. Our observations indicate elevated repair levels at domain boundaries with increased insulation, indicating these regions as potential repair hotspots. Conversely, domains with low insulation or lost boundaries exhibit nominal repair, with repair levels negatively correlated with domain length. We also noted chromatin loops displaying the highest strength appear to correlate with lower repair levels. Additionally, we categorized genomic bins based on their chromatin folding dynamics post-UV exposure, revealing a distinct relationship between local compaction/decompaction of the 3D genome and repair efficiency. Regions exhibiting high local decompaction (Comparison 3) demonstrated higher repair levels, while those with local compaction (Comparison 2) displayed lower repair levels. Furthermore, Comparison 1 identified regions acting as repair hubs, particularly for CPD repair. These hubs signify segments of the genome that come into close spatial proximity, facilitating open accessibility for damage recognition within the 3D genome context. Notably, as known within the NER machinery, CPDs and 6-4PPs exhibit different recognition and repair rates. While CPDs are poorly recognized by XPC alone, 6-4PPs are efficiently recognized by XPC-RAD23B, a crucial initiator of global-genome nucleotide excision repair [63]. Our observations at the 12-minute mark post-UV exposure highlight the divergence in damage recognition between CPDs and 6-4PPs, with the 6-4PPs displaying a more disoriented structure conducive to efficient recognition. Given these insights, focusing on CPD repair offers a more convenient framework for elucidating the facilitative roles of the 3D genome within the proposed model of decompaction and damage recognition.

After UV-induced DNA damage, besides facilitating efficient DNA repair, the genome must reorganize in accordance with transcriptional regulation upon genotoxic stress. Accordingly, UV radiation can impact cellular responses through the activation of various signaling pathways involving immediate early genes like AP-1 (Activator Protein-1) subunits, such as *JUN* and *FOS* family. For instance, AP-1 is important for inflammation induced by tumor necrosis factor alpha (TNF-α) signalling, which we found as the most significant hallmark in the time-course expression analysis. *JUN* and *FOS* protects cells from UV-induced cell death and cooperates with NF-κB to prevent apoptosis induced by TNF-α [64].

Both *JUN* and *FOS* are involved in regulating cell cycle progression and apoptosis following UV exposure. One key finding is that, loop anchors that lost interaction strength post-UV signifies the potential binding of transcription factor JUN and FOS. This was also accompanied by SMAD3 that is also important for TFG-beta signalling in the context of AP-1. In the GNN analysis, Comparison 2 might suggest that post-UV related gain of contact intensity at these regions might be regulated with loop extrusion, leading to denser and more strict structures. On the other hand, central regions of Comparison 3 showed highly significant enrichment for FOS, JUN-family proteins along with FOSL-2, and ATF3, suggesting potential AP-1 activity that is well-known after UV radiation [20,65]. On the other hand, we see NR3C1 TF enrichment at Comparison 3 sites, while its expression is downregulated. This potentially shows lower binding at Comparison 3 sites, and limited effect on AP-1 transrepression [66,67].

Overall, our study sheds light on the intricate interplay between UV-induced DNA damage, higher-order chromatin organization, and transcriptional regulation. We demonstrate that UV radiation induces significant alterations across various levels of chromatin organization, reminiscent of responses seen after ionizing radiation. Notably, we observe changes in distal interactions, compartmentalization dynamics, and TAD boundaries, which may serve as indicators of the cellular response to genotoxic stress. Our findings suggest that the 3D genome architecture plays a crucial role in facilitating efficient DNA repair, particularly through the modulation of accessibility for repair machinery. Furthermore, our investigation into the role of AP-1 subunits, such as JUN and FOS, highlights their involvement in orchestrating cellular responses to UV radiation. Moving forward, our research will focus on generating additional genomic datasets, including JUN/FOS and CTCF ChIP-seq post-UV, to further validate and refine our proposed model. By integrating these datasets with our existing data, we aim to gain deeper insights into the intricate mechanisms underlying higher-order chromatin reorganization and the DNA damage response.

## Conclusion

Our study represents the first comprehensive analysis of a unique multi-omics dataset, examining how the 3D genome orchestrates the response to UV-induced DNA damage. Specifically, we focused on the earliest time points within the first hour following UV exposure to follow immediate effects.

Using a multi-omics integrative approach, we investigated deeply sequenced Hi-C and RNA-seq datasets, aligning them with XR-Seq and Damage-Seq datasets from our previous study [24]. We conducted a comprehensive analysis, employing hierarchical and exploratory methods to investigate genome architectures. Additionally, we utilized a deep learning approach to uncover the mechanisms underlying UV-exposed genome folding. Moreover, we explored the intricate interplay among 3D chromatin structure, transcriptional regulation, and DNA damage and repair processes. This comprehensive approach shed light on the pivotal role of genome folding in the immediate response to UV damage.

## Methods

### Cell culture and exposure of human cells to UV

HeLa-S3 cells were maintained in Dulbeco’s modified Eagle’s medium (PAN-P04-03500) supplemented with 10% (v/v) heat-inactivated fetal bovine serum (PAN-P30-3304), 100 U/ml penicillin/streptomycin (PAN-P06-07100), 2 mM L-glutamine (PAN-P04-80100) and 1x MEM non-essential amino acid (Gibco 11140-35) solution. Cells were maintained at 37 °C with 5% CO2. Cells were grown to 80% confluence in 150-mm tissue culture dishes. UV treatment has been applied as described. [9] Briefly, culture medium was first discarded, and 15 ml of 1× PBS (PAN-P04-036503) per dish were used to wash the cells and discarded. With the cover off, cells were placed under a germicidal lamp emitting 254-nm UV (1 J/m2/s) for 20 s, then added 20 ml of DMEM to the tissue culture dish, and incubated the cells at 37°C for the recovery time points of 12, 30 and 60 minutes. Digital UVX Radiometer (UVP 97001601) was used to ensure that the cells were exposed to the desired UV dosage, and for reproducibility. Cells were immediately followed by Hi-C or RNA-Seq library preperation.

### Hi-C Library Preparation and Sequencing

Hi-C library was generated using the Phase Genomics Proximo Human Kit version 4.0. Cells were cross-linked for 15 minutes at room temperature with end-over-end mixing. Crosslinking reaction was terminated with quenching solution for 20 minutes at room temperature again with end-over-end mixing. Quenched cells were rinsed once with 1X Chromatin Rinse Buffer (CRB). A low-speed spin was used to remove the supernatant and the pellet containing the nuclear fraction of the lysate was washed with 1X CRB. After removing 1X CRB wash, the pellet was resuspended in 100 µL Proximo Lysis Buffer 2 and incubated at 65°C for 15 minutes. Chromatin was bound to Recovery Beads for 10 minutes at room temperature, placed on a magnetic stand, and washed with 200 µL of 1X CRB. Chromatin bound on beads was resuspended in 150 µL of Proximo fragmentation mix and incubated for 1 hour at 37°C. Reaction was cooled to 12°C and incubated with 2.5 µL of finishing enzyme for 30 minutes. Following the addition of 6 µL of Stop Solution, the beads were washed with 1X CRB and resuspended in 95 µL of Proximo Ligation Buffer supplemented with 5 µL of Proximity ligation enzyme. Proximity ligation reaction was incubated at room temperature for 4 hours with end-over-end mixing. To this volume, 5 µL of Reverse Crosslinks enzyme was added and the reaction incubated at 65°C for 1 hour. After reversing crosslinks, the free DNA was purified with Recovery Beads and Hi-C junctions were bound to streptavidin beads and washed to remove unbound DNA. Washed beads were used to prepare paired-end deep sequencing libraries using the Proximo Library preparation reagents. The libraries were sequenced on Illumina NovaSeq 6000 platform with PE150 reads.

### Hi-C Data Preprocessing

After preprocessing and quality control of FASTQ files with fastp[68], raw sequencing reads were aligned to human genome (GRCh38, Gencode v35). The data was processed using the Juicer v.1.6 pipeline (Supp. Table1). [69] Only chr1:22 and chrX were processed, and proceeded to downstream analysis. We used Juicer’s calculate_map_resolution.sh script to determine the maximum data resolution for each sample, finding that 10kb is the highest resolution available across all samples. This resolution was determined based on the criterion that the number of bins with more than 1000 contacts should constitute at least 80% of the total number of bins. Accordingly, .hic files that were generated with Juicer have been converted .mcool format with hicConvertFormat subprogram of HiCExplorer to be utilized for downstream analysis. [70] To make samples comparable and remove systematic bias, contact matrices were iteratively corrected with ICE [71] algorithm implementation with balance subprogram within cooler version 0.9.2[72], while keeping only cis-interactions (intra-chromosomal), filtering bins that are in blacklisted regions [73], ignoring first 2 diagonals, and using variance convergence threshold of 1e-05. We have obtained distance-dependent decay of interaction counts at 10kb resolution for each Hi-C sample, using “expected_cis” function of cooltools version 0.5.4 [74], and focused on interactions up to a linear distance of 10mb. We have utilized powerlaw version 1.5.0 [75] to obtain decay parameters for the hyper-tailed distrubutions.

### Compartment Analysis

To obtain eigenvectors for compartment profiles using eigen decomposition, we have utilized “eigs_cis” function of cooltools version 0.5.4 [74], at 100kb matrix resolution. For phasing between A and B profiles, we have tested POLR2A TF ChIP-seq (ENCSR000BGO), RNA-Seq, and GC content to assign bins to correct compartments, in which higher correlation associates to correct eigenvector for compartments. We have generated merged RNA-Seq tracks for each sample using “bamCoverage” function in deepTools version 3.5.4 [76] using BPM normalization, in order to test correlation for eigenvectors. With all of the phasing tracks, principal eigenvector were the one with highest correlation among first three eigenvectors (Supp. Table2).

Accordingly, 100kb bins were grouped into 50 percentiles based on 1st eigenvector. Saddle plots have been calculated with “saddle” function of cooltools considering the observed interactions normalized by expected, in which top left corners corresponds to B compartments, while bottom right corners correspond to A compartments. We have quantified the compartment strength using the “saddle_strength” function of cooltools, which computes the ratio between (AA+BB) and (AB+BA), by comparing the observed/expected values. With respect to saddle plots, compartment strength corresponds to comparing the ratio of values in the upper left and lower right corners to those in the lower left and upper right corners in the plot with an increasing extent. Thus, compartmentalization strength is calculated by assessing the relative enrichment of interactions within the same compartment (AA+BB) compared to interactions between different compartments (AB+BA). Accordingly, we have used this metric to compare degree of segregation between active and inactive chromatin regions between Hi-C samples.

In order to analyze compartment correlations between timepoints with respect to RNA-Seq data, we have obtained 1st eigenvectors for each Hi-C sample and assigned 100kb bins to factors of A1, A0, B0 and B1. Accordingly, for each gene in each cluster that we have obtained from clustering analysis, we have overlapped the compartment factors (100kb bins) to associated genic regions using “overlap” function in bioframe version 0.4.1 [77]. For each pair of sample correlation given the cluster, we have performed polychoric correlation with “polychoric_correlation” function in RyStats version 0.5.0 with respect to compartment factors. To perform bootstrapping, we have randomly picked 100kb bins depending on the number of bins on the original overlap to genes for each cluster and sample-pair comparison, 1000 times. Accordingly, we have compared original correlation to the correlation values in the bootstrapped distrubution (expected correlation) to generate two-tailed p-values.

### Boundary and TADs Analysis

In order to identify boundaries to see how genome is segmented into distinct domains, we have utilized “insulation” function in cooltools version 0.5.4 [74]. Briefly, insulation profiles have been calculated for each bin at 10kb resolution using the sliding window with the size of 500kb. Insulation profiles indicate specific sites with reduced scores, indicating decreased contact frequencies between adjacent genomic regions, and often these sites represent boundaries [78]. Accordingly, strongly insulating regions have been identified as boundaries, and boundary strength is calculated using peak prominence method on the minima of the insulation profiles.

To obtain preserved and non-preserved boundaries between each timepoint transition, we have identified boundaries for each Hi-C sample. Boundaries were considered to be preserved with a tolerated range of ± 10kb. Subsequently, by merging consecutive boundaries and excluding excessively long domains (>1.5mb), we have compiled a set of coordinates delineating topologically associating domains (TADs). Contact matrix snippets for pile-up analysis (aka. Aggregate Peak Analysis) plots were piled-up with coolpuppy version 1.1.0 [79] for boundaries on-diagonal.

We have calculated domain scores (TAD strength) as described [43]. Briefly, a domain score is the ratio of within-TAD intensity to between-TAD intensity on the rescaled contact matrix snippets spanning each TAD and flanking regions. Rescaled contact matrix snippets for TADs have been obtained with coolpuppy version 1.1.0 [79].

### Chromatin Loop Analysis

Chromatin loops were identified using the “dots” function within cooltools version 0.5.4 [74], operating at a resolution of 10kb. In summary, default kernels were applied as per cooltools recommendations for the resolution, which included the donut kernel as described in Rao et. al [26], along with vertical and horizontal kernels to mitigate the identification of stripes as loops, and a lower-left kernel to prevent the misidentification of pixels at domain corners as loops. Loops with a false discovery rate (FDR) value greater than 0.1, adjusted for multiple hypothesis testing using the Benjamini-Hochberg method, were filtered out, retaining only significantly enriched pixels for each sample. During each transition between time points, we identified loops and their associated anchors that are shared within a permissible range of ± 50kb, categorizing them as common loops or anchors. Accordingly, loops falling outside this range are deemed specific to a particular time point in the transition, suggesting potential enrichment or loss of interaction strength relative to transition.

We have obtained CTCF binding sites using the Bioconducter package CTCF [80], and utilized “ctcf_orientation.py” accessible under <u>CTCF_orientation</u> to obtain CTCF orientation on the loop anchors. Also, we have utilized “LoopWidth_piechart.py” accessible under <u>LoopWidth</u> to obtain categorized loop lengths.

Off-diagonal snippets of the loops were piled-up with coolpuppy version 1.1.0 [79], with a flank of 100kb, generating a 21×21 matrix, in which central pixel corresponds to the interaction of the loop anchors. Corners scores of off-diagonal pile-ups are calculated with the ratio of the central pixel intensity to the mean pixel intensity of the 6×6 matrix on the associated corner (aka. Peak to Corner scores). To calculate loop strength scores for an individual loop, we have used the ratio of central pixel intensity to the mean intensity of remaining pixels, and assigned a loop score to each individual loop (aka. Peak to Mean score).

Genome tracks of Hi-C, RNA-Seq, CTCF levels and related histone markers are visualized with Figeno version 1.0.5 [81]. CTCF, H3K27me3, H3K4me3, H3K4me1, H3K27ac, and H3K9ac data were obtained from ENCODE [82] data portal with the following identifiers: ENCSR000AOA, ENCSR000APB, ENCSR000AOF, ENCSR000APW, ENCSR000AOC, and ENCSR000AOH.

### GNN Worklows for Detecting Regions Centering Topology Change from Contact Maps

For training phase of the graph neural network (GNN) model in order to learn a subgraph-isomorphism violation landscape for Hi-C contact differentiations, we have preprocessed megamap at 10kb resolution of Hi-C experiments done on HeLa-S3 with merged replicates (4DN data portal: 4DNESCMX7L58) [83,84]. Briefly, we have iteratively corrected the contact matrix with ICE[71] algorithm implementation with balance subprogram within cooler version 0.9.2[72], as we described in Hi-C data preprocessing. Accordingly, we have obtained observed over expected matrices per chromosome arm using “_make_cis_obsexp_fetcher” function in cooltools version 0.5.4 [74].

Followingly, we have preprocessed Hi-C contacts per matrix using quantiles information with “QuantileTransformer” function in Scikit-learn version 1.3.0 into 1000 quantiles with Uniform distrubution, with the aim of making different datasets comparable, thus transforming contacts into 0 to 1 range.

To generate a graph dataset for training, we randomly selected a bin and constructed a graph from the 51×51 matrix where the selected bin is the central bin, corresponding to 250kb flanking regions at a resolution of 10kb, totalling to 510kb regions. We have decided on such linear distance to understand local topology change in 3D, in accordance to distance-dependent decay comparisons between non-UV to post-UV 12 minutes sample in which short-to-mid range interactions are particularly favored post-UV. We iterated through this process until the total number of generated graphs reached 10% of the total available bins per matrix. Furthermore, we have encoded node features in a graph into two categories; central node or non-central node, and relative linear distances scaled on the distance to central node to ensure non-central nodes have unique identities. Accordingly, transformed Hi-C contacts were assigned as the edge features.

For the training process, we constructed positive and negative target/query pairs as mini-batches, while ensuring central nodes are preserved. Positive pairs were generated by randomly selecting a graph and sampling a target graph from the available bins, followed by sampling a query graph from the selected target graph. Conversely, negative pairs were generated by randomly selecting two graphs, ensuring that they are central to different genomic bins. A target graph was then sampled from one of the selected graphs, while a query graph was sampled from the other ensuring number of nodes are smaller. By employing a target graph sampling strategy that encompasses a varying number of nodes within the range of (31, 51], we aim to train our model to generalize more effectively for contact maps with varying/nominal interaction coverage. This methodology enables the model to adeptly adapt to various graph sizes, enhancing its capability to navigate scenarios where contact maps might lack extensive coverage, while disregarding areas with sparse interactions.

We opted for a model design featuring 3 GNN layers, a choice informed by the observation that the diameter of query graphs approximates 3 in our Hi-C samples. This decision ensures that the model’s architecture aligns well with the inherent structure and complexity of the graphs, facilitating effective information propagation and feature extraction across the network. For the choice of GNN operator, we adopted Graph Isomorphism Network (GIN) [85] with a slight modification described by Hu et al. [86], in order to incorporate edge features into message-passing between nodes, referred as “GINE”. We chose to adopt the subgraph prediction function for training from the Neuromatch [58] algorithm because of its demonstrated capability to capture subgraph relations in the embedding space by enforcing the order embedding constraint. Briefly, the function evaluates whether a target graph node’s embedding has a k-hop neighborhood that is subgraph isomorphic to the k-hop neighborhood of a center node in the query graph, based on the embeddings of these nodes. It achieves this by minimizing the max margin loss, ensuring that positive pairs exhibit embeddings where the query node’s elements are less than or equal to the corresponding elements in the target node’s embedding. Conversely, for negative pairs, the violation of the subgraph constraint must exceed a certain margin in order to have zero loss. Furthermore, we also propagate amount of violation of target and query pairs to a classifier, referred to as the comparator, with the aim of learning a threshold for distinguishing between “Low” and “High” scores.

Therefore, we aimed to compare non-UV and 12min Hi-C samples (10kb bin resolution), considering this as the initial transition after UV-exposure and a reference point aligned with the timepoint of XR-Seq, to learn about altering topologies and immediate early response. To infer, we applied identical preprocessing steps to both of these contact maps as those used for the 4DN megamap, also disregarding blacklisted, balance-masked and sparsely interacting regions. Accordingly, diverging from the graph dataset generation process during the training phase, we have generated target and query pairs using a sliding window approach, with the strict number of node constraints, tolerating maximum node number of 41, and 51 for query and target, respectively. Therefore, we have acquired violation scores for each 10kb bin genome-wide, and learned the labels for distinguishing between “High” and “Low” scores.

GNN model has been implemented with Pytorch version 2.0.1 [87] and Pytorch Geometric version 2.3.1 [88]. Training and inference workflows were implemented with Pytorch Lightning version 2.0.6 in order to optimize distributed data parallel strategy for multi-processing. The model has been trained on Nvidia A100 in Sabanci University’s ToSUN HPC with Slurm Workload Manager.

### Transcription-factor differential enrichment analysis

We have performed differential enrichment analysis for TF binding using Unibind workflow [49]. Unibind is a database containing experimentally derived transcription factor binding sites (TFBSs) specific to various cell types or tissues, obtained from ChIP-seq experiments. Briefly, we have obtained Unibind’s LOLA database hg38_robust_UniBind_LOLA.RDS v3, from the corresponding Zenodo repository. In order to perform differential enrichment, we have preprocessed genomic coordinates in bed format, and ran LOLA version 1.32.0 [89] for enrichment calculation and statistical significance. We have only kept the experiment sets that their enrichments are significant (p-value < 0.05). And also, we have only kept the transcription factors that are supported by at least 10 individual experiment sets.

### RNA-Seq Library Preparation and Sequencing

RNA isolation was carried out using Genezol RNA isolation reagent (Geneaid GZR100) following the manufacturer’s instructions. A total amount of 0.4 μg RNA per sample was used as input material for the RNA sample preparations. Sequencing libraries were generated using NEBNext® UltraTM RNA Library Prep Kit for Illumina® (NEB, USA) following manufacturer’s recommendations and index codes were added to attribute sequences to each sample. Briefly, mRNA was purified from total RNA using poly-T oligo-attached magnetic beads. Fragmentation was carried out using divalent cations under elevated temperature in NEBNext First Strand Synthesis Reaction Buffer (5X). First strand cDNA was synthesized using random hexamer primer and M-MuLV Reverse Transcriptase (RNase H-). Second strand cDNA synthesis was subsequently performed using DNA Polymerase I and RNase H. The remaining overhangs were converted into blunt ends via exonuclease/polymerase activities. After adenylation of 3’ ends of DNA fragments, NEBNext Adaptor with hairpin loop structure were ligated to prepare for hybridization. In order to select cDNA fragments of preferentially 250∼300bp in length, the library fragments were purified with AMPure XP system (Beckman Coulter, Beverly, USA). Then PCR was performed with Phusion High-Fidelity DNA polymerase, Universal PCR primers and Index (X) Primer. At last, PCR products were purified (AMPure XP system) and library quality was assessed on the Agilent Bioanalyzer 2100 system.

The clustering of the index-coded samples was performed on a cBot Cluster Generation System using TruSeq PE Cluster Kit v3-cBot-HS (Illumia) according to the manufacturer’s instructions. After cluster generation, the libraries were sequenced on an Illumina Novaseq 6000 platform and 150bp paired-end reads were generated.

### RNA-Seq Data Preprocessing

After preprocessing and quality control of FASTQ files with fastp[68], raw sequencing reads have been aligned to human genome (GRCh38, Gencode v35) and its corresponding annotation file, with STAR aligner version 2.7.6a [doi:10/f4h523] with arguments –genomeLoad NoSharedMemory –quantMode TranscriptomeSAM –twopassMode Basic –outSAMtype BAM SortedByCoordinate. Followingly, transcripts (transcripts per million) were quantified using Salmon version 1.6.0[90].

### Gene Expression and Time-Series Analysis

Transcript-level abundance, estimated counts and transcript lengths, were imported with Bioconducter package tximport version 1.28.0[91] to be utilized for downstream analysis. Differential gene expression analysis was conducted with Bioconducter package DESeq2 version 1.40.2 [92]. Pre-filtering was applied to the data sets to filter low read counts in which genes with low number of reads were removed from the analysis. The read counts across the samples were normalised using the DESeq method, based on median ratio of gene counts. In order to perform time series analysis, likelihood ratio test was utilized with the full model on the time point factor, and the reduced model on the intercept. Resulting genes were filtered statistically (adjusted p-value ≤ 0.05) to obtain differentially expressed genes over time.

### Clustering and Gene Set Over-Representation Analysis

Clustering of the genes with respect to time-series analysis was performed with Bioconducter package coseq version 1.24.0[35]. DESeq normalized counts were used as input to coseq, without further normalization within coseq internally. Arcsine transformation was applied to expression matrix, and co-expression analysis of the differentially expressed genes were performed with Normal mixture model. Genes that were uniquely assigned to clusters with maximum conditional probability (min: 0.9) were retained for cluster visualization and over-representation analysis per each cluster.

Statistically enriched (adjusted p-value ≤ 0.05) terms of Gene Ontology Biological Process [36] and The Molecular Signatures DB Hallmarks [37], on genes assigned to each co-expression cluster were identified using Bioconducter package clusterProfiler version 4.8.1.[93] Over-representation analysis for each cluster was based on the background of the total diffentially expressed genes with respect to time factor, obtained from the likelihood ratio test. For gene sets created in the chromatin loop and GNN comparison analysis in Python, we have used “enrichr” function in GSEApy Python package version 1.0.6 [94] to perform over-representation analysis while also using NCI-Nature PID database [62].

### XR-Seq and Damage-Seq Data Preprocessing and Simulation

Raw XR-Seq and Damage-Seq reads were accessed under SRA PRJNA608124. We have processed the raw reads with XR-Seq and Damage-Seq Snakemake workflow version 0.7, that is accessible under <u>xr-</u> <u>ds-seq-snakemake</u>[24], with Snakemake version 7.32.4 [95]. Briefly, adapter sequences (TGGAATTCTCGGGTGCCAAGGAACTCCAGTNNNNNNACGATCTCGTATGCCGTCTTCTGCTTG) were trimmed from 3′ ends of raw XR-seq reads, and adapter sequences (GACTGGTTCCAATTGAAAGTGCTCTTCCGATCT) were trimmed from 5′ ends of raw Damage-seq reads using cutadapt version 4.1 [96]. Then, trimmed reads were aligned to human genome GRCh38 using Bowtie2 version 2.4.1 [97]. Resulting bam files were converted to bed using bedtools version 2.29.0 [98]. Also, aligned reads were sorted and duplicated regions were removed. Accordingly, we have processed the XR-Seq and Damage-Seq data for the exact repair and damage site coordinates, mapped to 1kb intervals on human genome, and performed Counts Per Million (CPM) normalization.

We have also generated synthetic NGS reads from the input DNA sequencing of HeLa-S3 (SRA:PRJNA608124) using NGS read simulator Boquila version 0.6.0 [99]. Briefly, the simulation tool Boquila randomly picks genomic regions to ensure that the chosen pseudo-reads match the nucleotide frequency of the provided NGS dataset. Accordingly, we have generated synthetic XR-Seq and Damage-Seq datasets, and utilized in order to have an expected background for repair and damage maps genome-wide with respect to dimer frequency. We have also performed CPM normalization for synthetic reads on 1kb intervals. Normalized “Repair levels” and “Damage sensitivity” are calculated with the normalization of original XR-Seq and Damage-Seq datasets to their correponding synthetic datasets. Furthermore, “Repair efficiency” is calculated using “Repair levels” normalized with “Damage sensitivity”.

### Software

We conducted our analyses using the Python language version 3.11.0 and R language version 4.3.1 with Bioconductor version 3.17. The manuscript was written and edited collaboratively on GitHub using Manubot.[100] Plots were generated with seaborn version 0.13.0[101], and figures were arranged on the open-source vector graphics software Inkscape.

## Acknowledgements

We would like to thank Dr. Ozlem Kutlu (Sabancı University Nanotechnology Research and Application Center, Türkiye) for invaluable support by providing access to the lab facilities for us to facilitate cell culture and sample preparation. We thank Adebali Lab members, Dr. Marat Kazanov (Faculty of Engineering and Natural Sciences, Sabanci University, Istanbul, Turkey), and Dr. Ferhat Ay (Center for Autoimmunity and Inflammation, La Jolla Institute for Immunology, La Jolla, CA, 92037, USA) for their feedbacks on the manuscript.

## Data Availability

All raw sequencing data generated in this study have been submitted to the <u>NCBI SRA database</u>, under accession number <u>PR</u>J<u>NA1053630</u>. Processed files in order to reproduce the analysis code were also made available on GEO database. Multi-resolution contact matrices in mcool format can be found with the identifier <u>GSE268350</u> for Hi-C data, and TPM quantification data can be found with the identifier <u>GSE268349</u> for RNA-seq data.

## Code Availability

All of the genome architecture analysis code and, research code for dataset generation, training, inference for GNN algorithm have been deposited at Github <u>CompGenomeLab/uv-3d-ddr</u>.

## Ethics declarations

### Competing interests

The authors declare no competing interests.

## Contributions

OA and VOK conceived of the project and designed the experiments. VOK performed cell culture, sample preparation and computational analyses. VOK, and OA interpreted the data and wrote the first draft. All authors have read, revised and approved the final manuscript.

## Notes

### Competing Interest Statement

The authors have declared no competing interest.

## References

1. Formation of UV-induced DNA damage contributing to skin cancer development Jean Cadet, Thierry Douki Photochemical & Photobiological Sciences (2018-12) grdgkb DOI: 10.1039/c7pp00395a · PMID: 29405222

2. RELATIVE INDUCTION OF CYCLOBUTANE DIMERS and CYTOSINE PHOTOHYDRATES IN DNA IRRADIATED in vitro and in vivo WITH ULTRAVIOLET-C and ULTRAVIOLET-B LIGHT David L Mitchell, Jin Jen, James E Cleaver Photochemistry and Photobiology (1991-11) cgp92d DOI: 10.1111/j.1751-1097.1991.tb02084.x · PMID: 1665910

3. Molecular Mechanisms of Ultraviolet Radiation-Induced DNA Damage and Repair Rajesh P Rastogi, Richa, Ashok Kumar, Madhu B Tyagi, Rajeshwar P Sinha Journal of Nucleic Acids (2010) bcjsd4 DOI: 10.4061/2010/592980 · PMID: 21209706 · PMCID: PMC3010660

4. Cell cycle arrest and apoptosis provoked by UV radiation-induced DNA damage are transcriptionally highly divergent responses M Gentile Nucleic Acids Research (2003-08-15) d7jthr DOI: 10.1093/nar/gkg675 · PMID: 12907719 · PMCID: PMC169943

5. UV Damage and DNA Repair in Malignant Melanoma and Nonmelanoma Skin Cancer Knuth Rass, Jörg Reichrath Sunlight, Vitamin D and Skin Cancer dsj4wc DOI: 10.1007/978-0-387-77574-6_13 · PMID: 18348455

6. Review: Ultraviolet radiation and skin cancer Deevya L Narayanan, Rao N Saladi, Joshua L Fox International Journal of Dermatology (2010-08-30) fjfstm DOI: 10.1111/j.1365-4632.2010.04474.x · PMID: 20883261

7. Mechanisms of DNA Repair by Photolyase and Excision Nuclease (Nobel Lecture) Aziz Sancar Angewandte Chemie International Edition (2016-06-23) f3qhrr DOI: 10.1002/anie.201601524 · PMID: 27337655

8. Nucleotide Excision Repair in Mammalian Cells Richard D Wood Journal of Biological Chemistry (1997-09) fxf9vg DOI: 10.1074/jbc.272.38.23465 · PMID: 9295277

9. Genome-wide mapping of nucleotide excision repair with XR-seq Jinchuan Hu, Wentao Li, Ogun Adebali, Yanyan Yang, Onur Oztas, Christopher P Selby, Aziz Sancar Nature Protocols (2018-12-14) ggkvth DOI: 10.1038/s41596-018-0093-7 · PMID: 30552409 · PMCID: PMC6429938

10. Dynamic maps of UV damage formation and repair for the human genome Jinchuan Hu, Ogun Adebali, Sheera Adar, Aziz Sancar Proceedings of the National Academy of Sciences (2017-06-12) ggdfws DOI: 10.1073/pnas.1706522114 · PMID: 28607063 · PMCID: PMC5495279

11. Loop extrusion as a mechanism for formation of DNA damage repair foci Coline Arnould, Vincent Rocher, Anne-Laure Finoux, Thomas Clouaire, Kevin Li, Felix Zhou, Pierre Caron, PhilippeE Mangeot, Emiliano P Ricci, Raphaël Mourad, … Gaëlle Legube Nature (2021-02-17) gnbt3b DOI: 10.1038/s41586-021-03193-z · PMID: 33597753 · PMCID: PMC7116834

12. Double-strand break repair and mis-repair in 3D Jennifer Zagelbaum, Jean Gautier DNA Repair (2023-01) gtkz8b DOI: 10.1016/j.dnarep.2022.103430 · PMID: 36436496 · PMCID: PMC10799305

13. Genome-wide mapping of long-range contacts unveils clustering of DNA double-strand breaks at damaged active genes François Aymard, Marion Aguirrebengoa, Emmanuelle Guillou, Biola M Javierre, Beatrix Bugler, Coline Arnould, Vincent Rocher, Jason S Iacovoni, Anna Biernacka, Magdalena Skrzypczak, … Gaëlle Legube Nature Structural & Molecular Biology (2017-03-06) f9s42d DOI: 10.1038/nsmb.3387 · PMID: 28263325 · PMCID: PMC5385132

14. Multiscale reorganization of the genome following DNA damage facilitates chromosome translocations via nuclear actin polymerization Jennifer Zagelbaum, Allana Schooley, Junfei Zhao, Benjamin R Schrank, Elsa Callen, Shan Zha, Max E Gottesman, André Nussenzweig, Raul Rabadan, Job Dekker, Jean Gautier Nature Structural & Molecular Biology (2022-12-23) gtkz8c DOI: 10.1038/s41594-022-00893-6 · PMID: 36564591 · PMCID: PMC10104780

15. Chromatin Compaction Protects Genomic DNA from Radiation Damage Hideaki Takata, Tomo Hanafusa, Toshiaki Mori, Mari Shimura, Yutaka Iida, Kenichi Ishikawa, Kenichi Yoshikawa, Yuko Yoshikawa, Kazuhiro Maeshima PLoS ONE (2013-10-09) gtk2bj DOI: 10.1371/journal.pone.0075622 · PMID: 24130727 · PMCID: PMC3794047

16. Radiation-induced DNA damage and repair effects on 3D genome organization Jacob T Sanders, Trevor F Freeman, Yang Xu, Rosela Golloshi, Mary A Stallard, Ashtyn M Hill, Rebeca San Martin, Adayabalam S Balajee, Rachel Patton McCord Nature Communications (2020-12-02) gshj9d DOI: 10.1038/s41467-020-20047-w · PMID: 33268790 · PMCID: PMC7710719

17. DNA double-strand breaks induce H2Ax phosphorylation domains in a contact-dependent manner Patrick L Collins, Caitlin Purman, Sofia I Porter, Vincent Nganga, Ankita Saini, Katharina E Hayer, Greer L Gurewitz, Barry P Sleckman, Jeffrey J Bednarski, Craig H Bassing, Eugene M Oltz Nature Communications (2020-06-22) gmzjq8 DOI: 10.1038/s41467-020-16926-x · PMID: 32572033 · PMCID: PMC7308414

18. ATM signaling modulates cohesin behavior in meiotic prophase and proliferating cells Zhouliang Yu, Hyung Jun Kim, Abby F Dernburg Nature Structural & Molecular Biology (2023-03-06) gr3whk DOI: 10.1038/s41594-023-00929-5 · PMID: 36879153 · PMCID: PMC10113158

19. Chromatin compartmentalization regulates the response to DNA damage Coline Arnould, Vincent Rocher, Florian Saur, Aldo S Bader, Fernando Muzzopappa, Sarah Collins, Emma Lesage, Benjamin Le Bozec, Nadine Puget, Thomas Clouaire, … Gaëlle Legube Nature (2023-10-18) gtgxsv DOI: 10.1038/s41586-023-06635-y · PMID: 37853125 · PMCID: PMC10620078

20. The mammalian UV response: mechanism of DNA damage induced gene expression. P Herrlich, C Sachsenmaier, A Radler-Pohl, S Gebel, C Blattner, HJ Rahmsdorf Advances in enzyme regulation (1994) https://www.ncbi.nlm.nih.gov/pubmed/7942283 DOI: 10.1016/0065-2571(94)90024-8 · PMID: 7942283

21. Ultraviolet B Regulation of Transcription Factor Families: Roles of Nuclear Factor-kappa B (NF-&#954;B) and Activator Protein-1 (AP-1) in UVB-Induced Skin Carcinogenesis S Cooper, G Bowden Current Cancer Drug Targets (2007-06-01) fgw37f DOI: 10.2174/156800907780809714 · PMID: 17979627 · PMCID: PMC2605645

22. Differential Regulation of the AP-1 Family Members by UV Irradiation In Vitro and In Vivo Kirsi Isoherranen, Jukka Westermarck, Veli-Matti Kähäri, Christer Jansén, Kari Punnonen Cellular Signalling (1998-03) cj4pjj DOI: 10.1016/s0898-6568(97)00100-9 · PMID: 9607142

23. The Mammalian UV Response Eitan Shaulian, Martin Schreiber, Fabrice Piu, Michelle Beeche, Erwin F Wagner, Michael Karin Cell (2000-12) fg6g64 DOI: 10.1016/s0092-8674(00)00193-8 · PMID: 11136975

24. Effects of replication domains on genome-wide UV-induced DNA damage and repair Yanchao Huang, Cem Azgari, Mengdie Yin, Yi-Ying Chiou, Laura A Lindsey-Boltz, Aziz Sancar, Jinchuan Hu, Ogun Adebali PLOS Genetics (2022-09-26) gtgsfn DOI: 10.1371/journal.pgen.1010426 · PMID: 36155646 · PMCID: PMC9536635

25. Comprehensive Mapping of Long-Range Interactions Reveals Folding Principles of the Human Genome Erez Lieberman-Aiden, Nynke L van Berkum, Louise Williams, Maxim Imakaev, Tobias Ragoczy, Agnes Telling, Ido Amit, Bryan R Lajoie, Peter J Sabo, Michael O Dorschner, … Job Dekker Science (2009-10-09) fnh876 DOI: 10.1126/science.1181369 · PMID: 19815776 · PMCID: PMC2858594

26. A 3D Map of the Human Genome at Kilobase Resolution Reveals Principles of Chromatin Looping Suhas SP Rao, Miriam H Huntley, Neva C Durand, Elena K Stamenova, Ivan D Bochkov, James T Robinson, Adrian L Sanborn, Ido Machol, Arina D Omer, Eric S Lander, Erez Lieberman Aiden Cell (2014-12) xqj DOI: 10.1016/j.cell.2014.11.021 · PMID: 25497547 · PMCID: PMC5635824

27. Physical mechanisms behind the large scale features of chromatin organization Ana Pombo, Mario Nicodemi Transcription (2014-03-21) grpsdn DOI: 10.4161/trns.28447 · PMID: 25764220 · PMCID: PMC4214237

28. Liquid chromatin Hi-C characterizes compartment-dependent chromatin interaction dynamics Houda Belaghzal, Tyler Borrman, Andrew D Stephens, Denis L Lafontaine, Sergey V Venev, Zhiping Weng, John F Marko, Job Dekker Nature Genetics (2021-02-11) gjh93f DOI: 10.1038/s41588-021-00784-4 · PMID: 33574602 · PMCID: PMC7946813

29. Chromatin architecture reorganization during stem cell differentiation Jesse R Dixon, Inkyung Jung, Siddarth Selvaraj, Yin Shen, Jessica E Antosiewicz-Bourget, Ah Young Lee, Zhen Ye, Audrey Kim, Nisha Rajagopal, Wei Xie, … Bing Ren Nature (2015-02-18) f6zxg5 DOI: 10.1038/nature14222 · PMID: 25693564 · PMCID: PMC4515363

30. Deciphering aging at three-dimensional genomic resolution Zunpeng Liu, Juan Carlos Izpisua Belmonte, Weiqi Zhang, Jing Qu, Guang-Hui Liu Cell Insight (2022-06) gtktmv DOI: 10.1016/j.cellin.2022.100034 · PMID: 37193050 · PMCID: PMC10120299

31. Reorganization of chromosome architecture in replicative cellular senescence Steven W Criscione, Marco De Cecco, Benjamin Siranosian, Yue Zhang, Jill A Kreiling, John M Sedivy, Nicola Neretti Science Advances (2016-02-05) gf8b6q DOI: 10.1126/sciadv.1500882 · PMID: 26989773 · PMCID: PMC4788486

32. Dynamic 3D genome reorganization during senescence: defining cell states through chromatin Haitham A Shaban, Susan M Gasser Cell Death & Differentiation (2023-08-18) gtktmx DOI: 10.1038/s41418-023-01197-y · PMID: 37596440

33. Rapid reversible changes in compartments and local chromatin organization revealed by hyperosmotic shock Ramon Amat, René Böttcher, François Le Dily, Enrique Vidal, Javier Quilez, Yasmina Cuartero, Miguel Beato, Eulàlia de Nadal, Francesc Posas Genome Research (2018-12-06) gkgc8p DOI: 10.1101/gr.238527.118 · PMID: 30523037 · PMCID: PMC6314167

34. Genomic meta-analysis of the interplay between 3D chromatin organization and gene expression programs under basal and stress conditions Idan Nurick, Ron Shamir, Ran Elkon Epigenetics & Chromatin (2018-08-29) gtktmz DOI: 10.1186/s13072-018-0220-2 · PMID: 30157915 · PMCID: PMC6114837

35. Transformation and model choice for RNA-seq co-expression analysis Andrea Rau, Cathy Maugis-Rabusseau Briefings in Bioinformatics (2017-01-08) gdjvgk DOI: 10.1093/bib/bbw128 · PMID: 28065917

36. The Gene Ontology knowledgebase in 2023 Gene Ontology Consortium, Suzi A Aleksander, James Balhoff, Seth Carbon, JMichael Cherry, Harold J Drabkin, Dustin Ebert, Marc Feuermann, Pascale Gaudet, Nomi L Harris, … Monte Westerfield Genetics (2023-05-04) https://www.ncbi.nlm.nih.gov/pmc/articles/PMC10158837/ DOI: 10.1093/genetics/iyad031 · PMID: 36866529 · PMCID: PMC10158837

37. Molecular signatures database (MSigDB) 3.0 Arthur Liberzon, Aravind Subramanian, Reid Pinchback, Helga Thorvaldsdóttir, Pablo Tamayo, Jill P Mesirov Bioinformatics (2011-05-05) b8mx73 DOI: 10.1093/bioinformatics/btr260 · PMID: 21546393 · PMCID: PMC3106198

38. Three-dimensional genome architecture: players and mechanisms Ana Pombo, Niall Dillon Nature Reviews Molecular Cell Biology (2015-03-11) gfx6c7 DOI: 10.1038/nrm3965 · PMID: 25757416

39. Targeted Degradation of CTCF Decouples Local Insulation of Chromosome Domains from Genomic Compartmentalization Elphège P Nora, Anton Goloborodko, Anne-Laure Valton, Johan H Gibcus, Alec Uebersohn, Nezar Abdennur, Job Dekker, Leonid A Mirny, Benoit G Bruneau Cell (2017-05) f98xmm DOI: 10.1016/j.cell.2017.05.004 · PMID: 28525758 · PMCID: PMC5538188

40. Cohesin Loss Eliminates All Loop Domains Suhas SP Rao, Su-Chen Huang, Brian Glenn St Hilaire, Jesse M Engreitz, Elizabeth M Perez, Kyong-Rim Kieffer-Kwon, Adrian L Sanborn, Sarah E Johnstone, Gavin D Bascom, Ivan D Bochkov, … Erez Lieberman Aiden Cell (2017-10) gb4ggf DOI: 10.1016/j.cell.2017.09.026 · PMID: 28985562 · PMCID: PMC5846482

41. Two independent modes of chromatin organization revealed by cohesin removal Wibke Schwarzer, Nezar Abdennur, Anton Goloborodko, Aleksandra Pekowska, Geoffrey Fudenberg, Yann Loe-Mie, Nuno A Fonseca, Wolfgang Huber, Christian H Haering, Leonid Mirny, Francois Spitz Nature (2017-09-27) gfc4zv DOI: 10.1038/nature24281 · PMID: 29094699 · PMCID: PMC5687303

42. Functional Analysis of CTCF During Mammalian Limb Development Natalia Soshnikova, Thomas Montavon, Marion Leleu, Niels Galjart, Denis Duboule Developmental Cell (2010-12) ff7xxj DOI: 10.1016/j.devcel.2010.11.009 · PMID: 21145498

43. Single-nucleus Hi-C reveals unique chromatin reorganization at oocyte-to-zygote transition Ilya M Flyamer, Johanna Gassler, Maxim Imakaev, Hugo B Brandão, Sergey V Ulianov, Nezar Abdennur, Sergey V Razin, Leonid A Mirny, Kikuë Tachibana-Konwalski Nature (2017-03-29) f9v63d DOI: 10.1038/nature21711 · PMID: 28355183 · PMCID: PMC5639698

44. Molecular mechanisms and genomic maps of DNA excision repair in Escherichia coli and humans Jinchuan Hu, Christopher P Selby, Sheera Adar, Ogun Adebali, Aziz Sancar Journal of Biological Chemistry (2017-09) gftwfh DOI: 10.1074/jbc.r117.807453 · PMID: 28798238 · PMCID: PMC5612094

45. Coming full circle: On the origin and evolution of the looping model for enhancer– promoter communication Tessa M Popay, Jesse R Dixon Journal of Biological Chemistry (2022-08) gtktmw DOI: 10.1016/j.jbc.2022.102117 · PMID: 35691341 · PMCID: PMC9283939

46. Enhancer–promoter interactions and transcription are largely maintained upon acute loss of CTCF, cohesin, WAPL or YY1 Tsung-Han S Hsieh, Claudia Cattoglio, Elena Slobodyanyuk, Anders S Hansen, Xavier Darzacq, Robert Tjian Nature Genetics (2022-12) grxc8f DOI: 10.1038/s41588-022-01223-8 · PMID: 36471071 · PMCID: PMC9729117

47. Enhancer-Promoter Communication: It’s Not Just About Contact Annabelle Wurmser, Srinjan Basu Frontiers in Molecular Biosciences (2022-04-19) gtktm2 DOI: 10.3389/fmolb.2022.867303 · PMID: 35517868 · PMCID: PMC9061983

48. CTCF: making the right connections Rodolfo Ghirlando, Gary Felsenfeld Genes & Development (2016-04-15) f8kfqd DOI: 10.1101/gad.277863.116 · PMID: 27083996 · PMCID: PMC4840295

49. UniBind: maps of high-confidence direct TF-DNA interactions across nine species Rafael Riudavets Puig, Paul Boddie, Aziz Khan, Jaime Abraham Castro-Mondragon, Anthony Mathelier BMC Genomics (2021-06-26) gs9mk3 DOI: 10.1186/s12864-021-07760-6 · PMID: 34174819 · PMCID: PMC8236138

50. A joint NCBI and EMBL-EBI transcript set for clinical genomics and research Joannella Morales, Shashikant Pujar, Jane E Loveland, Alex Astashyn, Ruth Bennett, Andrew Berry, Eric Cox, Claire Davidson, Olga Ermolaeva, Catherine M Farrell, … Terence D Murphy Nature (2022-04-06) gqn657 DOI: 10.1038/s41586-022-04558-8 · PMID: 35388217 · PMCID: PMC9007741

51. Multiscale and integrative single-cell Hi-C analysis with Higashi Ruochi Zhang, Tianming Zhou, Jian Ma Nature Biotechnology (2021-10-11) gpg3tn DOI: 10.1038/s41587-021-01034-y · PMID: 34635838 · PMCID: PMC8843812

52. scHiCEmbed: Bin-Specific Embeddings of Single-Cell Hi-C Data Using Graph Auto-Encoders Tong Liu, Zheng Wang Genes (2022-06-11) grxcdt DOI: 10.3390/genes13061048 · PMID: 35741810 · PMCID: PMC9222580

53. Prediction of gene co-expression from chromatin contacts with graph attention network Ke Zhang, Chenxi Wang, Liping Sun, Jie Zheng Bioinformatics (2022-08-05) gtjtdv DOI: 10.1093/bioinformatics/btac535 · PMID: 35929807 · PMCID: PMC9525008

54. HiC-GNN: A generalizable model for 3D chromosome reconstruction using graph convolutional neural networks Van Hovenga, Jugal Kalita, Oluwatosin Oluwadare Computational and Structural Biotechnology Journal (2023) gszk9r DOI: 10.1016/j.csbj.2022.12.051 · PMID: 36698967 · PMCID: PMC9842867

55. Subgraph extraction and graph representation learning for single cell Hi-C imputation and clustering Jiahao Zheng, Yuedong Yang, Zhiming Dai Briefings in Bioinformatics (2023-11-22) gtjtdt DOI: 10.1093/bib/bbad379 · PMID: 38040494 · PMCID: PMC10691963

56. Improving comparative analyses of Hi-C data via contrastive self-supervised learning Han Li, Xuan He, Lawrence Kurowski, Ruotian Zhang, Dan Zhao, Jianyang Zeng Briefings in Bioinformatics (2023-06-07) gtjtds DOI: 10.1093/bib/bbad193 · PMID: 37287135

57. A graph neural network-based interpretable framework reveals a novel DNA fragility– associated chromatin structural unit Yu Sun, Xiang Xu, Lin Lin, Kang Xu, Yang Zheng, Chao Ren, Huan Tao, Xu Wang, Huan Zhao, Weiwei Tu, … Xiaochen Bo Genome Biology (2023-04-24) gtjtdw DOI: 10.1186/s13059-023-02916-x · PMID: 37095580 · PMCID: PMC10124043

58. Neural Subgraph Matching Rex, Ying, Zhaoyu Lou, Jiaxuan You, Chengtao Wen, Arquimedes Canedo, Jure Leskovec arXiv (2020) gtgv88 DOI: 10.48550/arxiv.2007.03092

59. Structural and Functional Properties of Activator Protein-1 in Cancer and Inlammation Pritam Bhagwan Bhosale, Hun Hwan Kim, Abuyaseer Abusaliya, Preethi Vetrivel, Sang Eun Ha, Min Yeong Park, Ho Jeong Lee, Gon Sup Kim Evidence-Based Complementary and Alternative Medicine (2022-05-26) gqtsxg DOI: 10.1155/2022/9797929 · PMID: 35664945 · PMCID: PMC9162854

60. AP-1: a double-edged sword in tumorigenesis Robert Eferl, Erwin F Wagner Nature Reviews Cancer (2003-11) b354hs DOI: 10.1038/nrc1209 · PMID: 14668816

61. The AP-1 transcriptional complex: Local switch or remote command? Fabienne Bejjani, Emilie Evanno, Kazem Zibara, Marc Piechaczyk, Isabelle Jariel-Encontre Biochimica et Biophysica Acta (BBA) - Reviews on Cancer (2019-08) gndg6f DOI: 10.1016/j.bbcan.2019.04.003 · PMID: 31034924

62. PID: the Pathway Interaction Database Carl F Schaefer, Kira Anthony, Shiva Krupa, Jeffrey Buchoff, Matthew Day, Timo Hannay, Kenneth H Buetow Nucleic Acids Research (2008-10-02) dv62wn DOI: 10.1093/nar/gkn653 · PMID: 18832364 · PMCID: PMC2686461

63. Structure and mechanism of pyrimidine–pyrimidone (6-4) photoproduct recognition by the Rad4/XPC nucleotide excision repair complex Debamita Paul, Hong Mu, Hong Zhao, Ouathek Ouerfelli, Philip D Jeffrey, Suse Broyde, Jung-Hyun Min Nucleic Acids Research (2019-05-20) gtmcjj DOI: 10.1093/nar/gkz359 · PMID: 31106376 · PMCID: PMC6614856

64. c-Jun regulates cell cycle progression and apoptosis by distinct mechanisms R Wisdom The EMBO Journal (1999-01-04) fj366v DOI: 10.1093/emboj/18.1.188 · PMID: 9878062 · PMCID: PMC1171114

65. Ultraviolet-A induces activation of AP-1 in cultured human keratinocytes. M Djavaheri-Mergny, JL Mergny, F Bertrand, R Santus, C Mazière, L Dubertret, JC Mazière FEBS letters (1996-04-08) https://www.ncbi.nlm.nih.gov/pubmed/8797811 DOI: 10.1016/0014-5793(96)00294-3 · PMID: 8797811

66. Selective modulation of the glucocorticoid receptor can distinguish between transrepression of NF-κB and AP-1 Karolien De Bosscher, Ilse M Beck, Lien Dejager, Nadia Bougarne, Anthoula Gaigneaux, Sébastien Chateauvieux, Dariusz Ratman, Marc Bracke, Jan Tavernier, Wim Vanden Berghe, … Guy Haegeman Cellular and Molecular Life Sciences (2013-06-20) f5nbq6 DOI: 10.1007/s00018-013-1367-4 · PMID: 23784308 · PMCID: PMC3889831

67. Repression of transcription by the glucocorticoid receptor: A parsimonious model for the genomics era Anthony N Gerber, Robert Newton, Sarah K Sasse Journal of Biological Chemistry (2021-01) gtgxkc DOI: 10.1016/j.jbc.2021.100687 · PMID: 33891947 · PMCID: PMC8141881

68. Ultrafast one-pass FASTQ data preprocessing, quality control, and deduplication using fastp Shifu Chen iMeta (2023-05) gs9rt7 DOI: 10.1002/imt2.107

69. Juicer Provides a One-Click System for Analyzing Loop-Resolution Hi-C Experiments Neva C Durand, Muhammad S Shamim, Ido Machol, Suhas SP Rao, Miriam H Huntley, Eric S Lander, Erez Lieberman Aiden Cell Systems (2016-07) gfc8h7 DOI: 10.1016/j.cels.2016.07.002 · PMID: 27467249 · PMCID: PMC5846465

70. Galaxy HiCExplorer 3: a web server for reproducible Hi-C, capture Hi-C and single-cell Hi-C data analysis, quality control and visualization Joachim Wolff, Leily Rabbani, Ralf Gilsbach, Gautier Richard, Thomas Manke, Rolf Backofen, Björn A Grüning Nucleic Acids Research (2020-04-17) ghjzhc DOI: 10.1093/nar/gkaa220 · PMID: 32301980 · PMCID: PMC7319437

71. Iterative correction of Hi-C data reveals hallmarks of chromosome organization Maxim Imakaev, Geoffrey Fudenberg, Rachel Patton McCord, Natalia Naumova, Anton Goloborodko, Bryan R Lajoie, Job Dekker, Leonid A Mirny Nature Methods (2012-09-02) gdcfmm DOI: 10.1038/nmeth.2148 · PMID: 22941365 · PMCID: PMC3816492

72. Cooler: scalable storage for Hi-C data and other genomically labeled arrays Nezar Abdennur, Leonid A Mirny Bioinformatics (2019-07-10) gf6crw DOI: 10.1093/bioinformatics/btz540 · PMID: 31290943 · PMCID: PMC8205516

73. The ENCODE Blacklist: Identification of Problematic Regions of the Genome Haley M Amemiya, Anshul Kundaje, Alan P Boyle Scientific Reports (2019-06-27) gf4jsb DOI: 10.1038/s41598-019-45839-z · PMID: 31249361 · PMCID: PMC6597582

74. Cooltools: enabling high-resolution Hi-C analysis in Python, Nezar Abdennur, Sameer Abraham, Geoffrey Fudenberg, Ilya M Flyamer, Aleksandra A Galitsyna, Anton Goloborodko, Maxim Imakaev, Betul A Oksuz, Sergey V Venev Cold Spring Harbor Laboratory (2022-11-01) gtgs7n DOI: 10.1101/2022.10.31.514564

75. powerlaw: A Python Package for Analysis of Heavy-Tailed Distributions Jeff Alstott, Ed Bullmore, Dietmar Plenz PLoS ONE (2014-01-29) gc4n4j DOI: 10.1371/journal.pone.0085777 · PMID: 24489671 · PMCID: PMC3906378

76. deepTools2: a next generation web server for deep-sequencing data analysis Fidel Ramírez, Devon P Ryan, Björn Grüning, Vivek Bhardwaj, Fabian Kilpert, Andreas S Richter, Steffen Heyne, Friederike Dündar, Thomas Manke Nucleic Acids Research (2016-04-13) f8v7nt DOI: 10.1093/nar/gkw257 · PMID: 27079975 · PMCID: PMC4987876

77. Bioframe: Operations on Genomic Intervals in Pandas Dataframes, Nezar Abdennur, Geoffrey Fudenberg, Ilya Flyamer, Aleksandra A Galitsyna, Anton Goloborodko, Maxim Imakaev, Sergey V Venev Cold Spring Harbor Laboratory (2022-02-19) gtgs7m DOI: 10.1101/2022.02.16.480748

78. Cohesin-dependent globules and heterochromatin shape 3D genome architecture in S. pombe Takeshi Mizuguchi, Geoffrey Fudenberg, Sameet Mehta, Jon-Matthew Belton, Nitika Taneja, Hernan Diego Folco, Peter FitzGerald, Job Dekker, Leonid Mirny, Jemima Barrowman, Shiv IS Grewal Nature (2014-10-12) f6spq7 DOI: 10.1038/nature13833 · PMID: 25307058 · PMCID: PMC4465753

79. *Coolpup.py:* versatile pile-up analysis of Hi-C data Ilya M Flyamer, Robert S Illingworth, Wendy A Bickmore Bioinformatics (2020-01-31) ghq7rj DOI: 10.1093/bioinformatics/btaa073 · PMID: 32003791 · PMCID: PMC7214034

80. CTCF: an R/bioconductor data package of human and mouse CTCF binding sites Mikhail G Dozmorov, Wancen Mu, Eric S Davis, Stuart Lee, Timothy J Triche Jr., Douglas H Phanstiel, Michael I Love Bioinformatics Advances (2022-01-01) gtgvbp DOI: 10.1093/bioadv/vbac097 · PMID: 36699364 · PMCID: PMC9793704

81. Figeno: multi-region genomic figures with long-read support Etienne Sollier, Jessica Heilmann, Clarissa Gerhäuser, Michael Scherer, Christoph Plass, Pavlo Lutsik Cold Spring Harbor Laboratory (2024-04-26) gtvxpx DOI: 10.1101/2024.04.22.590500

82. New developments on the Encyclopedia of DNA Elements (ENCODE) data portal Yunhai Luo, Benjamin C Hitz, Idan Gabdank, Jason A Hilton, Meenakshi S Kagda, Bonita Lam, Zachary Myers, Paul Sud, Jennifer Jou, Khine Lin, … JMichael Cherry Nucleic Acids Research (2019-11-12) gqrkfz DOI: 10.1093/nar/gkz1062 · PMID: 31713622 · PMCID: PMC7061942

83. The 4D nucleome project Job Dekker, Andrew S Belmont, Mitchell Guttman, Victor O Leshyk, John T Lis, Stavros Lomvardas, Leonid A Mirny, Clodagh C O’Shea, Peter J Park, … Sheng Zhong Nature (2017-09-14) gfkhp8 DOI: 10.1038/nature23884 · PMID: 28905911 · PMCID: PMC5617335

84. The 4D Nucleome Data Portal as a resource for searching and visualizing curated nucleomics data Sarah B Reiff, Andrew J Schroeder, Koray Kırlı, Andrea Cosolo, Clara Bakker, Luisa Mercado, Soohyun Lee, Alexander D Veit, Alexander K Balashov, Carl Vitzthum, … Peter J Park Nature Communications (2022-05-02) gtgv86 DOI: 10.1038/s41467-022-29697-4 · PMID: 35501320 · PMCID: PMC9061818

85. How Powerful are Graph Neural Networks? Keyulu Xu, Weihua Hu, Jure Leskovec, Stefanie Jegelka arXiv (2018) gq6fgc DOI: 10.48550/arxiv.1810.00826

86. Strategies for Pre-training Graph Neural Networks Weihua Hu, Bowen Liu, Joseph Gomes, Marinka Zitnik, Percy Liang, Vijay Pande, Jure Leskovec arXiv (2019) gtgv87 DOI: 10.48550/arxiv.1905.12265

87. PyTorch: An Imperative Style, High-Performance Deep Learning Library Adam Paszke, Sam Gross, Francisco Massa, Adam Lerer, James Bradbury, Gregory Chanan, Trevor Killeen, Zeming Lin, Natalia Gimelshein, Luca Antiga, … Soumith Chintala arXiv (2019) gpmnkd DOI: 10.48550/arxiv.1912.01703

88. Fast Graph Representation Learning with PyTorch Geometric Matthias Fey, Jan Eric Lenssen arXiv (2019) gq6gqp DOI: 10.48550/arxiv.1903.02428

89. LOLA: enrichment analysis for genomic region sets and regulatory elements in R and Bioconductor Nathan C Sheffield, Christoph Bock Bioinformatics (2015-10-27) f8c88x DOI: 10.1093/bioinformatics/btv612 · PMID: 26508757 · PMCID: PMC4743627

90. Salmon provides fast and bias-aware quantification of transcript expression Rob Patro, Geet Duggal, Michael I Love, Rafael A Irizarry, Carl Kingsford Nature Methods (2017-03-06) gcw9f5 DOI: 10.1038/nmeth.4197 · PMID: 28263959 · PMCID: PMC5600148

91. Differential analyses for RNA-seq: transcript-level estimates improve gene-level inferences Charlotte Soneson, Michael I Love, Mark D Robinson F1000Research (2015-12-30) gdtgw8 DOI: 10.12688/f1000research.7563.1 · PMID: 26925227 · PMCID: PMC4712774

92. Moderated estimation of fold change and dispersion for RNA-seq data with DESeq2 Michael I Love, Wolfgang Huber, Simon Anders Genome Biology (2014-12) gd3zvn DOI: 10.1186/s13059-014-0550-8 · PMID: 25516281 · PMCID: PMC4302049

93. clusterProfiler 4.0: A universal enrichment tool for interpreting omics data Tianzhi Wu, Erqiang Hu, Shuangbin Xu, Meijun Chen, Pingfan Guo, Zehan Dai, Tingze Feng, Lang Zhou, Wenli Tang, Li Zhan, … Guangchuang Yu The Innovation (2021-08) gmg747 DOI: 10.1016/j.xinn.2021.100141 · PMID: 34557778 · PMCID: PMC8454663

94. GSEApy: a comprehensive package for performing gene set enrichment analysis in Python Zhuoqing Fang, Xinyuan Liu, Gary Peltz Bioinformatics (2022-11-25) gtgw4d DOI: 10.1093/bioinformatics/btac757 · PMID: 36426870 · PMCID: PMC9805564

95. Sustainable data analysis with Snakemake Felix Mölder, Kim Philipp Jablonski, Brice Letcher, Michael B Hall, Christopher H Tomkins-Tinch, Vanessa Sochat, Jan Forster, Soohyun Lee, Sven O Twardziok, Alexander Kanitz, … Johannes Köster F1000Research (2021-04-19) gj76rq DOI: 10.12688/f1000research.29032.2 · PMID: 34035898 · PMCID: PMC8114187

96. Cutadapt removes adapter sequences from high-throughput sequencing reads Marcel Martin EMBnet.journal (2011-05-02) gdh7xt DOI: 10.14806/ej.17.1.200

97. Fast gapped-read alignment with Bowtie 2 Ben Langmead, Steven L Salzberg Nature Methods (2012-03-04) gd2xzn DOI: 10.1038/nmeth.1923 · PMID: 22388286 · PMCID: PMC3322381

98. BEDTools: a lexible suite of utilities for comparing genomic features Aaron R Quinlan, Ira M Hall Bioinformatics (2010-01-28) cmrms3 DOI: 10.1093/bioinformatics/btq033 · PMID: 20110278 · PMCID: PMC2832824

99. Boquila: NGS read simulator to eliminate read nucleotide bias in sequence analysis ÜMİT AKKÖSE, OGÜN ADEBALİ Turkish Journal of Biology (2023-01-01) gtgsqz DOI: 10.55730/1300-0152.2650 · PMID: 37529166 · PMCID: PMC10387831

100. Open collaborative writing with Manubot Daniel S Himmelstein, Vincent Rubinetti, David R Slochower, Dongbo Hu, Venkat S Malladi, Casey S Greene, Anthony Gitter PLOS Computational Biology (2019-06-24) c7np DOI: 10.1371/journal.pcbi.1007128 · PMID: 31233491 · PMCID: PMC6611653

101. seaborn: statistical data visualization Michael Waskom Journal of Open Source Software (2021-04-06) gjqn3g DOI: 10.21105/joss.03021

